# Specific redox and iron homeostasis responses in the root tip of Arabidopsis upon zinc excess

**DOI:** 10.1101/2024.08.29.610234

**Authors:** Noémie Thiébaut, Ludwig Richtmann, Manon Sarthou, Daniel P. Persson, Alok Ranjan, Marie Schloesser, Stéphanie Boutet, Lucas Rezende, Stephan Clemens, Nathalie Verbruggen, Marc Hanikenne

**Affiliations:** InBioS-PhytoSystems, Translational Plant Biology, University of Liège, B-4000 Liège, Belgium; Laboratory of Plant Physiology and Molecular Genetics, Université Libre de Bruxelles, B-1050 Brussels, Belgium; Department of Plant Physiology and Faculty of Life Sciences: Food, Nutrition and Health, University of Bayreuth, 95440 Bayreuth, Germany; Department of Plant and Environmental Sciences, University of Copenhagen, 1871 Frederiksberg, Denmark; Université Paris-Saclay, INRAE, AgroParisTech, Institute Jean-Pierre Bourgin for Plant Sciences (IJPB), 78000 Versailles, France; Hedera-22 SA, Boulevard du Rectorat 27b, B-4000 Liège, Belgium

**Keywords:** Zinc, zinc excess, Root Apical Meristem, specialized metabolism, iron deficiency, metal transporter, Laser Ablation ICP-MS, camalexin, *Arabidopsis thaliana*

## Abstract

- Zinc (Zn) excess negatively impacts primary root growth in Arabidopsis. Yet, the effects of Zn excess on specific growth processes in the root tip remain largely unexplored.
- Transcriptomics, ionomics and metabolomics were used to examine the specific impact of Zn excess on the root tip (RT) compared to the remaining root (RR).
- Zn excess exposure resulted in shortened root apical meristem and elongation zone, with differentiation initiating closer to the tip of the root. Zn accumulated at a lower concentration in the RT than in RR. This pattern was associated with lower expression of Zn homeostasis and Fe deficiency response genes.
- A distinct distribution of Zn and Fe in RT and RR was highlighted by Laser Ablation ICP-MS analysis.
- Specialized Trp-derived metabolism genes, typically associated with redox and biotic stress responses, were specifically up-regulated in the RT upon Zn excess, among those *Phytoalexin Deficient 3* (*PAD3*) encoding the last enzyme of camalexin synthesis. In roots of wild-type seedlings, camalexin concentration increased by 6-fold upon Zn excess and a *pad3* mutant displayed increased Zn sensitivity and an altered ionome.
- Our results indicate that distinct redox and iron homeostasis mechanisms are key elements of the response to Zn excess in the RT.

## Introduction

In plants, the root system is responsible for nutrient uptake and translocation to the shoot. Upon differentiation of distinct root cell layers, many features contribute to the regulation of nutrient absorption and translocation. Root hair length and density are conditionally adjustable traits, which increase external surface, hence nutrient absorption (Leitner *et al*., 2010; Cai & Ahmed, 2022; van Dijk *et al*., 2022). Root vacuolar storage, which increases with the presence of bigger vacuoles in differentiated cells, permits controlling nutrient radial transport towards the stele and therefore also buffers root-to-shoot nutrient translocation (Ricachenevsky *et al*., 2015; Clemens, 2019). Several apoplastic non-permeable barriers, as for instance the Casparian strips or complete suberinization of endodermal cell walls, force nutrients into the symplastic pathway, and therefore to transmembrane transport (Barberon *et al*., 2016; Somssich *et al*., 2016; Barberon, 2017; Vestenaa *et al*., 2024). These traits associated with cell differentiation enable the regulation of the uptake and radial transport of nutrients by locally controlling the presence, (sub-)localization or activity of transmembrane nutrient transporters or facilitators (Barberon *et al*., 2016; Castaings *et al*., 2016; Sinclair *et al*., 2018; Hanikenne *et al*., 2021; Spielmann *et al*., 2022). Stress, such as nutrient excess or deficiency, usually results in a modification of these traits to maintain, within homeostatic limits, appropriate nutrition (Sinclair & Krämer, 2012; Ricachenevsky *et al*., 2015; Clemens, 2019).

Root barriers and other traits are specific of differentiated tissues, hence the regulation of nutrient distribution to undifferentiated tissues may differ (Kondo *et al*., 2014; Barberon *et al*., 2016; Barberon, 2017; Zluhan-Martínez *et al*., 2021). Such tissues are mainly found in root tips (RT), where quiescence is maintained, and meristematic activity takes place together with cell elongation and differentiation onset. These processes, controlled by hormonal and redox balances, are the motor of root growth and are affected by several nutrient stresses or metal excesses, *e. g.* zinc (Zn) or cadmium (Cd) excesses (De Smet *et al*., 2015; Yuan & Huang, 2016; Bruno *et al*., 2017; Zhang *et al*., 2018; Wang *et al*., 2021; Zluhan-Martínez *et al*., 2021; Leonardo *et al*., 2021; van Dijk *et al*., 2022).

Zinc (Zn) is an essential plant micronutrient (Amini *et al*., 2022; Stanton *et al*., 2022; Thiébaut & Hanikenne, 2022; Lilay *et al*., 2024), estimated to be a cofactor for ∼3000 proteins in *Arabidopsis thaliana* (Arabidopsis) (Clemens, 2022). Nonetheless, it is well established that Zn excess is toxic (Fukao *et al*., 2011; Shanmugam *et al*., 2011), although the initial molecular events leading to Zn toxicity at the cellular level remain unclear. Zn binding to functional groups of proteins, such as sulfhydryl groups, is thought to affect the protein activity and/or structure. Zn ions in excess can also compete with other ions for uptake and/or binding to proteins (van Dijk *et al*., 2022). In Arabidopsis, Zn excess impairs shoot and root growth, induces chlorosis and iron (Fe) deficiency, as Zn and Fe share transporters and metal chelators (Fukao *et al*., 2011; Shanmugam *et al*., 2011; Charlier *et al*., 2015; Lešková *et al*., 2017; Scheepers *et al*., 2020). Fe supplementation during Zn excess mitigates chlorosis and partially rescues growth inhibition, highlighting the major contribution of Fe imbalance to Zn toxicity (Fukao *et al*., 2011; Shanmugam *et al*., 2011; Lešková *et al*., 2017). Finally, Zn excess was shown to alter root architecture, to reduce the size of mature root cells (Fukao *et al*., 2011), and to impact the auxin gradient which maintains meristematic activity in the root apical meristem (RAM) (Sofo *et al*., 2017; Zhang *et al*., 2018; Wang *et al*., 2021). RAM maintenance also requires an appropriate Fe supply and an optimal redox balance (Henriques *et al*., 2002; Tsukagoshi *et al*., 2010; Reyt *et al*., 2015), which are both influenced by the Zn status (Remans *et al*., 2012; Cuypers *et al*., 2016; Hanikenne *et al*., 2021).

The molecular responses to Zn excess in differentiated roots of Arabidopsis are documented. Zn excess is essentially managed by regulating its transport and chelation, which limits uptake, favors root sequestration and restricts translocation to the shoot (Desbrosses-Fonrouge *et al*., 2005; Kaur & Garg, 2021), and by mitigating its secondary effects, such as Fe deficiency (Shanmugam *et al*., 2011; Hanikenne *et al*., 2021; Stanton *et al*., 2023). It mobilizes (i) several Zn transporter families, among which Zinc-regulated Iron-regulated transporter-like Proteins (ZIP), Heavy-Metal ATPase (HMA) or Metal Tolerance Proteins (MTP), involved in Zn uptake, vacuolar sequestration and/or translocation to the shoot (Sinclair & Krämer, 2012; Ricachenevsky *et al*., 2015; Kaur & Garg, 2021); (ii) metal ion chelators, like nicotianamine (NA), a Zn chelator contributing to vacuolar Zn retention upon excess (Haydon *et al*., 2012), or phytochelatins (PC) that are activated under Zn excess, increasing Zn chelation (Tennstedt *et al*., 2009; Clemens *et al*., 2013; Clemens, 2019); (iii) cell wall modifications, which influence cytoplasmic entry of excess metals, in particularly through pectins, whose modifications can increase the metal-binding capacity of the apoplast (Zhong *et al*., 2024).

Whereas the impact of Zn excess on root growth and root system architecture is well studied, the response mechanisms themselves in the RT remain to be investigated (van Dijk *et al*., 2022). Similarly, while well described in differentiated roots, Zn transport and distribution in the RT is yet to be explored. Here, we assessed the impact of Zn excess in the Arabidopsis RT, using a multi-omics approach. We showed that the ionome and transcriptome of the RT and differentiated roots were distinctly affected, suggesting specific regulation of Zn uptake and storage, as well as the Fe deficiency response in the RT. Using Laser Ablation Inductively Coupled Plasma-Mass Spectrometry (LA-ICP-MS), we highlighted differences in Fe and Zn localization between RT and differentiated roots. Finally, we showed that Trp-derived specialized metabolism was specifically triggered in the RT upon Zn excess.

## Materials and Methods

Extended method descriptions are provided in the Methods S1 text, as supporting information.

### Plant material and growth conditions

*A. thaliana* (Col-0) was used in all experiments. The *pad3-1* (CS3805), *pad3-3* (SALK_026585C), *pad3-4* (SALKseq_059784.1) mutants were obtained from NASC (Nottingham Arabidopsis Stock Centre) and used as homozygous lines.

Plants were grown vertically on agar plates with Hoagland medium (Scheepers *et al*., 2020) in BrightBoy GroBanks (CLF Plant Climatics) under long-days (16h light, 21°C, 100 µmol.m^-2^.s^-1^ / 8h dark, 18°C). For the LA-ICP-MS experiment, seedlings were grown under slightly different conditions [16h light, 22°C, 125 µmol.m^-2^.s^-1^ / 8h dark, 20°C, in a Percival chamber (CLF Plant Climatics)].

For most experiments, sterilized seeds were sown on control agar plates on nylon meshes and stratified (4°C, 2 days). After 7 days, seedlings were transferred to either new control (1 µM Zn) or Zn excess plates (200 µM Zn), and treated for 24 hours, 48 hours or 7 days (Fig. **S1**). For *pad3* mutant characterization, seeds were sown directly on control (1 µM Zn) or treated (75 µM Zn) plates, supplemented or not with 250 nM camalexin (Sigma Aldrich), stratified and grown for 11 days. Details of experimental replication levels are presented in each figure.

### Root growth and RAM phenotyping

Root growth was measured from scanned plate images after 24h, 48h and 7 days of treatment. Confocal microscope images of propidium iodide-stained roots were used to quantify meristematic zone length (Perilli & Sabatini, 2010), RAM cortex cell number and size, and elongation zone length. All features were measured using FIJI (https://imagej.net/Fiji).

### Ionome profiling

For each RT and RR sample, seedlings from 48 plates, each containing 16-18 seedlings, were pooled, washed and desorbed (Scheepers *et al*., 2020). About 2-3 mm long root tips (RT) were then separated from the remaining root system (RR). Dry samples were acid-digested under pressure using a high-performance microwave. Elemental analysis was performed using ICP-MS.

For the *pad3* mutant characterization, 11-day-old seedlings were harvested. Shoots and roots (60-90 seedlings per sample) were separated, washed, desorbed, and dried. Samples were acid-digested (Scheepers *et al*., 2020) and analyzed by ICP-OES (Inductively Coupled Plasma-Optical Emission Spectroscopy).

### Laser Ablation ICP-MS

Roots were harvested and two regions of interest (ROI) along the primary roots were prepared: RAM (∼200 µm from the tip of the columella) and differentiated roots (∼mid-distance from the apex) (Fig. **S2**). Following encapsulation in melted paraffin, 14 µm-thick transversal cryosections were prepared (Persson *et al*., 2016), then dried and analyzed using a LA-ICP-MS system.

### RNA-Seq analysis

For RNA sequencing, 48 plates (16-18 seedlings/plate) were harvested per independent replicate. Once separated, RT and RR samples were rapidly collected and frozen in liquid nitrogen (Fig. **S1**). Upon total RNA isolation, RNA quality, library preparation and quality control, and Illumina sequencing were performed on a NovaSeq6000 (Illumina) at the GIGA Genomics platform (ULiège, Belgium). Data quality assessment and trimming, read mapping, counting and differential expression analysis were performed as described (Scheepers *et al*., 2020). GO enrichment analysis for biological processes was conducted with the Panther Database.

### RT qPCR

Total RNA extraction, cDNA preparation, quantitative PCR, data normalization and analysis were performed as described (Scheepers *et al*., 2020). Primers are listed in Table **S1.** The *UBIQUITIN10* (*UBQ10*), *EF1alpha* and *AT1G58050* genes were used as references.

### Untargeted metabolomics analysis

For untargeted metabolomic analysis, 48h after the transfer to control or Zn excess plates, samples were taken by cutting and collecting similarly-sized pieces of RR and RT. Polar/semipolar primary and specialized metabolites were extracted and analyzed as described (Boutet *et al*., 2022). Relative metabolite quantifications were normalized to the total metabolite count of each sample.

### Targeted camalexin analysis

Roots and shoots from 11 day-old seedlings were separated and camalexin was extracted using a modified protocol from Tewes *et al*. (2018). Extracts were analyzed using Ultra-Performance Liquid Chromatography-High-Resolution Mass Spectrometry. Camalexin was identified based on the retention time of an external standard of commercial camalexin (Sigma Aldrich), and quantified using a calibration curve.

## Results

### Zn excess inhibits root growth and impacts the root apical meristem

One-week-old Arabidopsis seedlings exposed to control (1 µM Zn) or Zn excess (200 µM Zn) conditions showed noticeable primary root growth inhibition as early as 24h later, and after 48h and one week (Fig. **1a-b**). This phenotype was associated with a significant reduction of RAM size, both in micrometers and cell number after 48h, as observed in the cortex cell layer (Fig. **1c-e**). While cortex cell size was maintained in the division zone (delimited by red arrows in Fig. **1c**, Fig. **1f**), the elongation zone was shortened, and cell differentiation initiated closer to the tip of the root (Fig. **1g**). Indeed, the distance from the tip of the root to the first appearing root hair decreased by 1 and 1.5 mm after 24h and 48h of treatment, respectively (Fig. **1g**). Thus, root growth reduction upon Zn excess was associated with a reduction in size of both RAM and elongation zone.

**Figure 1.**
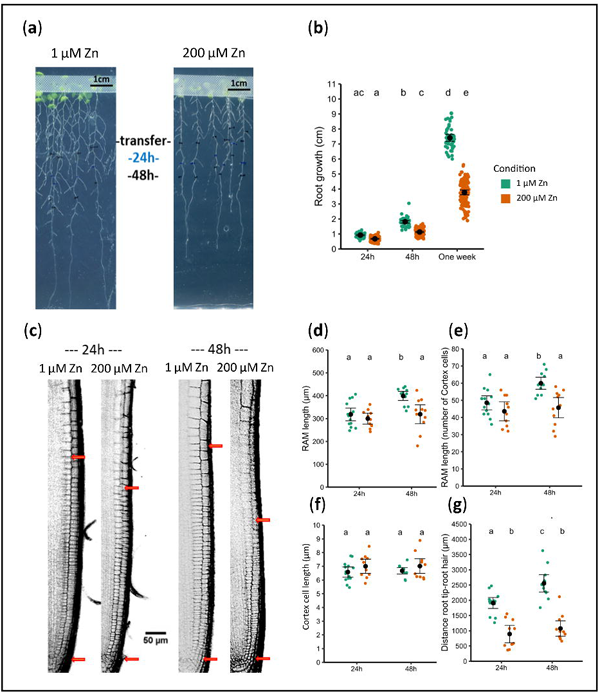
Primary root growth inhibition and impact on the root apical meristem upon Zn excess in Arabidopsis. Seedlings were germinated and grown for one week in control agar plates (1 µM Zn), then transferred for another week onto new control or Zn excess (200 µM Zn) plates. a. Representative pictures of seedlings one week after transfer. Marks on the plates represent the size of the primary roots at transfer, and after 24h and 48h. b. Primary root growth after transfer. Values are from 3 biological replicates each including 36 seedlings. c. Representative images of root apical meristem (RAM) 24h and 48h after transfer. RAM size was measured between the two red arrows pointing the quiescent center (QC) (lower arrow) and the first elongated cortex cell (upper arrow). d-g. RAM and elongation zone sizes 24h and 48h after transfer. RAM length was assessed as described in c, both in µm (d) and number of cells in the cortex cell layer (e). The mean cortex cell size in the RAM was obtained by dividing the length of RAM by the number of cortex cells for each meristem (f). First root hair distance was measured from the QC (g). d-g. Values are from 10 plants per condition per time-points. In b, d-g, black dots and whiskers represent mean values and standard deviations, respectively, whereas green (1 µM Zn) and orange (200 µM Zn) dots represent all individual values. Different letters correspond to significantly different groups (ANOVA type I, with Tukey test correction, *p-value* < 0.05).

### Zn excess differently impacts the root and root tip ionomes

As the root meristematic zone was already strongly affected by Zn excess after 24h and 48h, the ionome of the root tip (RT, ∼2-3 apical mm of primary root) and the remaining root system (RR) was examined at these time-points. Zn concentration dramatically increased in both RT and RR after 24h and 48h of exposure to Zn excess (Fig. **2a**). However, Zn accumulation was significantly lower in RT than in RR. Overall, the concentrations of other elements, including iron (Fe), remained mostly stable compared to control conditions. Exceptions were manganese (Mn) and magnesium (Mg), which decreased in RR after 48h of Zn excess (Fig. **2b**, Fig. **S3**) but not in RT. In contrast to most elements, phosphorus (P) concentrations were ∼3.5-fold higher in RT compared to RR in both control and Zn excess conditions. Conversely, Mn concentrations were markedly lower in RT than RR in control conditions (Fig. **2c**).

**Figure 2.**
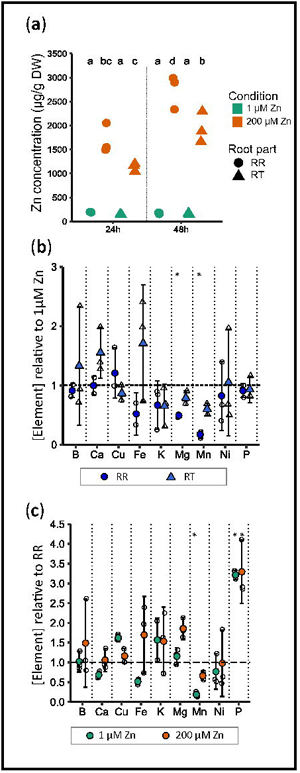
Ionome profiling of the root tip and the remaining root system upon Zn excess in Arabidopsis. Seedlings were germinated and grown for one week in control agar plates (1 µM Zn), then transferred for 24h or 48h onto new control or Zn excess (200 µM Zn) plates. Root tip (RT) and remaining root (RR) samples were analyzed by ICP-MS. a. Zn concentration. b. Element concentrations in the 200 µM Zn condition relative to their concentrations in control condition, 48h after transfer. c. Element concentrations in RT relative to their concentrations in RR, 48h after transfer. a-c. Values are from 3 independent replicates. Each replicate consisted of a pool of dissected tissues from 750-870 seedlings. Individual values (a) or shapes and whiskers representing mean values and standard deviations (b-c) are presented. Different letters correspond to significantly different groups (a) and stars represent statistical difference with ratio =1 (b-c) (ANOVA type I, with Tukey test correction, *p-value* < 0.05).

### Zn and Fe distributions in tissues differ between RR and RT

To examine the tissue-specific Zn distribution beyond bulk element concentrations, LA-ICP-MS analyses were conducted on sections across RAM and differentiated root tissues from seedlings grown in control and 48h Zn excess conditions. Fe distribution was additionally assessed in these samples, as Zn and Fe homeostasis are known to interact closely (Hanikenne *et al*., 2021). In control conditions, Zn was mostly present in the vascular cylinder of differentiated roots (Fig. **3a,c**), while in the RAM, the distribution was more uniform, with a stronger accumulation in the meristematic cell layer destined to differentiate into epidermis (Fig. **3b,d**). Overall, the Zn signal was slightly higher in the RAM (between ∼ 50 and 300 counts) than in differentiated roots (∼ 25-100 counts) in control conditions (Fig. **3c-d**). Conversely, Fe was preferentially localized in the epidermis of differentiated roots, while it concentrated in meristem cells destined to form the cortical ring (Fig. **3a-b**). Element distribution was thus distinct according to the metal (Zn vs Fe) and the root part (RAM vs differentiated tissue). Zn excess did not alter Zn and Fe distribution patterns, despite a strong increase in Zn accumulation in the tissues (Fig. **3a-d**).

**Figure 3.**
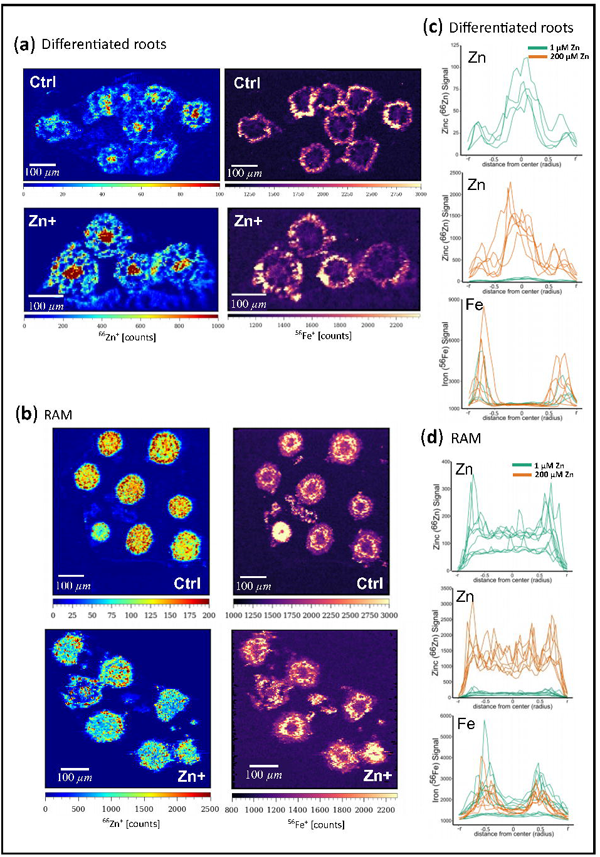
Zn and Fe distribution in the RAM and differentiated roots upon Zn excess in Arabidopsis. Seedlings were germinated and grown for one week in control agar plates (1 µM Zn), then transferred for 48h onto new control (Ctrl) or Zn excess (200 µM Zn, Zn+) plates. RAM and differentiated root transversal sections were then analyzed by Laser Ablation ICP-MS. a-b. ^66^Zinc and ^56^Fe imaging in differentiated roots (a) and RAM (b). c-d. Graphical representation of ^66^Zinc and ^56^Fe signals across the differentiated root (c) and RAM (d) cell layers. Each graph contains data for 8 root-or RAM-sections per condition, coming from two distinct root bouquets (see Materials and Methods and Fig. S2).

### RR and RT display distinct transcriptomic responses to Zn excess

RT and RR samples from seedlings exposed to either control or Zn excess conditions for 24h or 48h were submitted to RNA-Seq analysis. Principal component analysis (PCA) of the gene expression dataset showed a strong separation of RT and RR samples (PC1 explaining >95% of the transcriptional variance, Fig. **4a**), highlighting major differences between the two transcriptional landscapes. In contrast, treatment effects along PC2 only explained 2% of variance, and sampling time (24 or 48h) had little impact.

**Figure 4.**
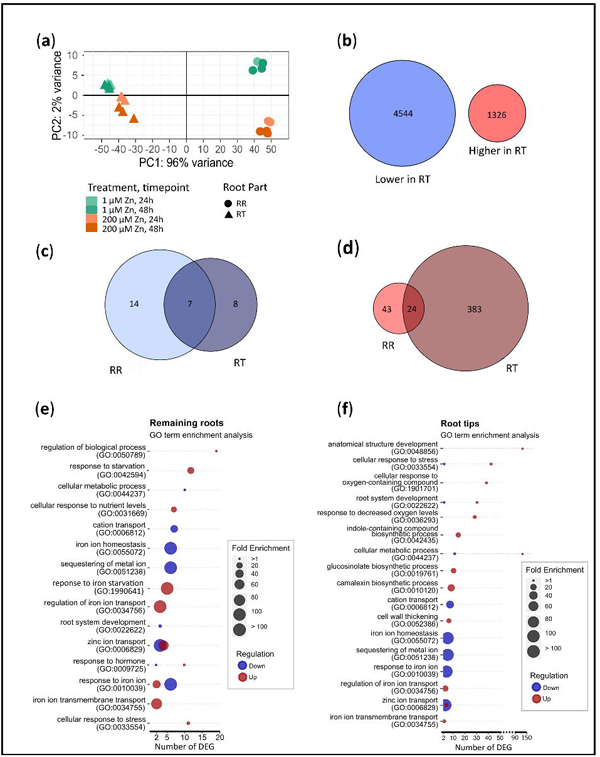
Transcriptome profiling of the root tip and the remaining root system upon Zn excess in Arabidopsis. Seedlings were germinated and grown for one week in control agar plates (1 µM Zn), then transferred for 24h or 48h onto new control or Zn excess (200 µM Zn) plates. Root tip (RT) and remaining root (RR) samples were then submitted to RNA-Seq. **a.** Principal Component Analysis of the RNA-Seq data **b.** Number of genes that are less (blue) or more (red) expressed in RT than in RR in control conditions, combining the 24h and 48h timepoints. **c-d.** Number of genes that are down-(**c**) or up-regulated (**d**) by Zn excess, both in RR and in RT. **b-d**. Differentially expressed genes (DEG) between control and Zn excess conditions were identified with the following criteria: log_2_ (fold change) < -1 of > 1; adjusted *p-value* < 0.05. **e-f.** Representative selection of enriched GO terms among DEGs that are up-or down-regulated upon Zn excess in RR (**f**) and RT (**g**). A complete description of the GO enrichment analysis is presented in (Fig. **S4-7**, and Table **S4**).

Consistent with the PCA analysis, a large number of genes exhibited differential expression between RT and RR. The number of Differentially Expressed Genes (DEGs) amounted to 5,870 in control conditions alone (Fig. **4b**, Table **S2**), with 4,374 of these genes also differing between RT and RR upon Zn excess (Fig. **S4a**). These DEGs reflected the morphological and physiological differences between RT and RR and were analyzed in more details in the accompanying manuscript (Richtmann *et al*.).

Furthermore, a total of 479 DEGs were identified as being up- or down-regulated by Zn excess in RR and/or in RT (Fig. **4c-d**, Table **S3**). Of these, 407 genes were up-regulated in RT by Zn excess. Noticeably, almost all DEGs identified after 24h of Zn excess were also identified after 48h (Fig. **S4b-e**), though the total number of DEGs increased at 48h, but essentially affected the same biological processes (Fig. **S5** and **S6**).

Gene Ontology (GO) enrichment analysis was conducted on all up- and down-regulated DEGs (24h and 48h combined) in RT and RR (Fig. **4e-f**, Fig. **S7**). A majority of Zn-regulated genes in RR corresponded to GOs linked to stress response and homeostatic processes, especially related to metal ions. These GOs were over-represented both among the up- and down-regulated DEGs (e.g., zinc ion transport GO:0006829). Fe homeostasis-related GOs were strongly represented, indicating an active Fe deficiency response: the ‘response to iron starvation’ and ‘iron sequestration’/‘iron ion homeostasis’ GOs were enriched among up- and down-regulated DEGs, respectively (Fig. **4e**).

In RT, GOs related to stress response and homeostatic processes were also prevalent. Most of the metal ion homeostasis GO-associated genes were down-regulated under Zn excess, with the exception of the ‘regulation of iron ion transport’ and ‘response to iron starvation’ GOs. In addition, RT Zn excess-responsive genes displayed more specific changes, that were not found in RR, such as GOs corresponding to (i) development (*i.e.* anatomical structure and root system development), (ii) reaction oxygen species and hypoxia, and (iii) specialized metabolism (*e.g.* camalexin, glucosinolate and indole-containing compound synthesis, tryptophan catabolism) (Fig. **4e-f**, Table **S4**).

In short, RT responded more strongly to Zn excess than RR, with a greater number of up-regulated genes. As a result, more critical processes, including developmental, redox, and specialized metabolism pathways, seemed to be influenced in RT. Conversely, ion homeostasis processes were observed in the transcriptomic responses of both root parts.

### The developmental impact of Zn excess is reflected in the RT transcriptome

Zn excess strongly reduced root growth and the size of the RAM (Fig. **1**), suggesting an impact of Zn excess on cell division. However, this process was not identified in the GO analysis and none of 46 marker genes for different phases of the cell cycle, endocycle, and DNA damage response were differentially expressed upon Zn excess (Fig. **S8**).

Zn excess induced root hair formation closer to the root apex (Fig. **1**), a developmental impact that was reflected in the RT transcriptomic profiling: the RNA-Seq PCA showed that RT and RR samples were less divergent along PC1 in Zn excess than in control conditions (Fig. **4a**), while the GO enrichment analysis highlighted the up-regulation of developmental processes in RT upon Zn excess (Fig. **4f**). Furthermore, cross-referencing DEGs upon Zn excess in RT and RR to cell-type specific expression data in the Arabidopsis roots (Shahan *et al*., 2022) showed increased differentiation of several cell layers in RT upon Zn excess: numerous genes (61) associated to differentiating cells were up-regulated (dark red in Fig. **S9**, Table **S5_Sheet 1**). Genes associated with trichoblast elongation and differentiation were the most affected (22 genes). Similarly, 24 genes associated with increased trichoblast elongation and differentiation were enriched among the genes more highly expressed in RT than in RR under Zn excess (Table **S5_Sheet 2**).

Finally, DEGs upon Zn excess in RT or those more highly expressed in RT than in RR solely in Zn excess, were cross-referenced with a list of ∼640 genes potentially implicated in root hair development (Won *et al*., 2009) (Table **S5_Sheets 3**-**4**), matching 90 and 75 genes, respectively.

Altogether, ∼30% of the 407 DEGs (*i.e.* 125 genes) up-regulated in RT upon Zn excess and 90 distinct genes more highly expressed in RT than in RR specifically in Zn excess condition, were implicated in the elongation or development of different cell layers (Fig. **S9**, Table **S5_Summary Sheet**). This included numerous expansins and cell wall modification-associated genes [e.g. *XYLOGLUCAN ENDOTRANSGLUCOSYLASE/HYDROLASE* (*XTH*), as discussed in more details in Richtmann *et al*.], transcription factors, actors of hormonal signaling (*e.g.* auxin or ethylene responses) and nutrition-associated genes (Table **S5**). Thus, a consequent part of the genes up-regulated upon Zn excess in the RT corresponded to root differentiation-associated genes, reflecting the Zn-triggered shift in the positioning of the different zones in the RT.

### Ionomic alterations upon Zn excess are reflected in the transcriptomic response

Among Zn excess-regulated genes associated with ion transport, only a few were linked to Zn transport, some to phosphate homeostasis, and finally, many to Fe transport and mobilization (Table **S6**). Hierarchical clustering of known Zn and Fe homeostasis genes according to their expression upon Zn excess or between root parts (RT vs RR) (Fig. **5**) revealed that most genes were significantly less expressed in RT, in both control and Zn excess conditions (in blue in Groups B-E of Fig. **5**), with few exceptions (*i.e.* Group A containing *ZIP4*, *MTP1*, *FER2*, *FER3*). Within Group B, three known Zn-deficiency responsive ZIP transporter genes (*IRT3*, *ZIP3* and *ZIP5*) (Talke *et al*., 2006; Van De Mortel *et al*., 2006; Assunção *et al*., 2010) were down-regulated by Zn excess in both RR (all 3 genes) and RT (*ZIP3* excluded), suggesting a general shut-down of cellular Zn uptake.

**Figure 5.**
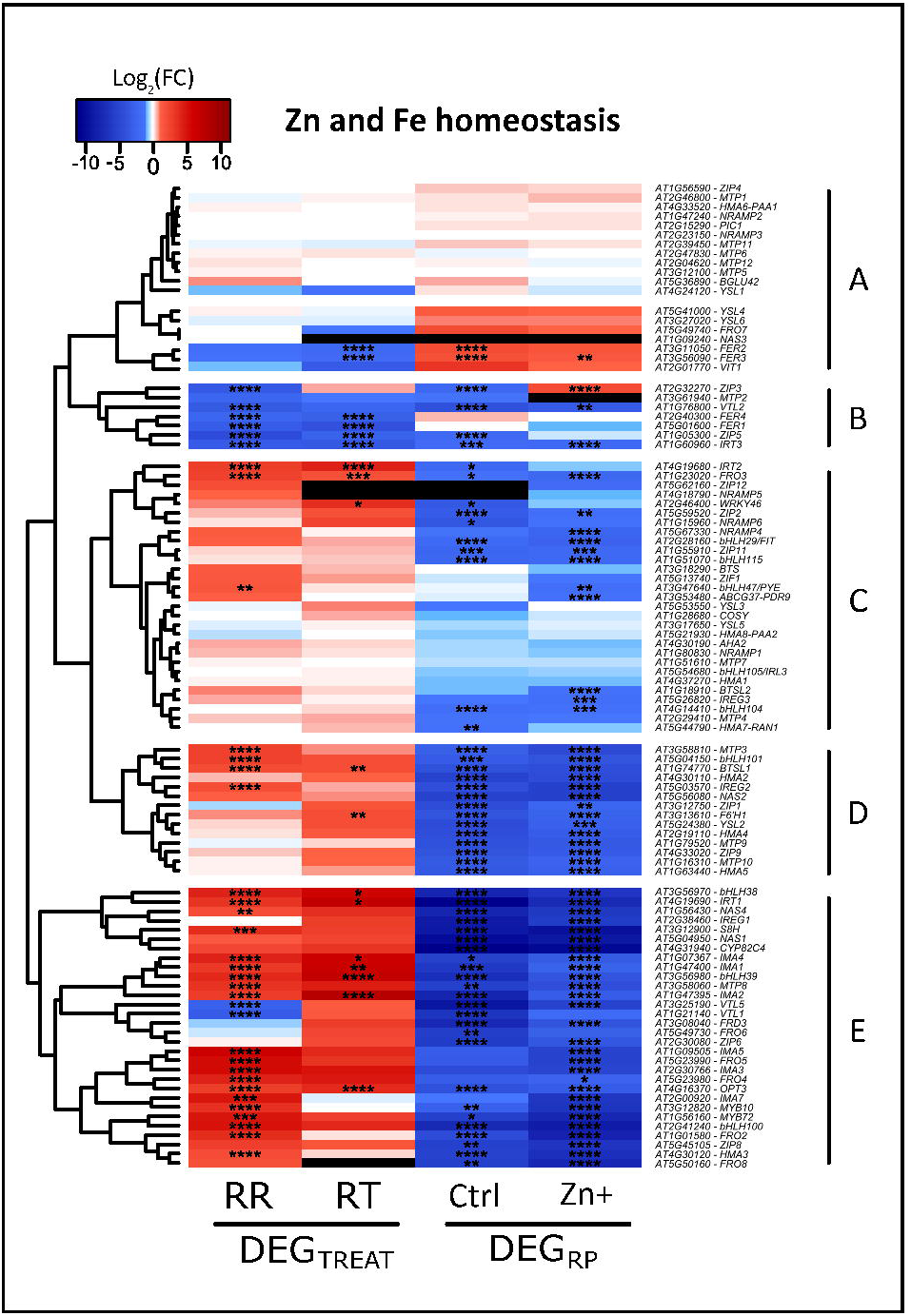
Expression of Zn and Fe homeostasis genes after 48h of Zn excess in Arabidopsis roots. Using the RNA-Seq data, the heatmap shows log_2_(fold change) of gene expression between Zn excess (Zn+) and control conditions (DEG_TREAT_) or between RT and RR (DEG_RP_) from a manually curated list of Zn and Fe homeostasis genes. Statistically-significant changes in gene expression (adjusted *p-value* < 0.05) are marked with stars:“ * ” for pval < 0.05, “ ** ” for pval <0.01, “ *** ” for pval < 0.001, “ **** ” for pval < 0.0001.

In contrast, *HMA3* and *MTP3* were strongly induced by Zn in RR (Fig. **5**, Groups D-E), suggesting increasing vacuolar Zn storage (Arrivault *et al*., 2006; Morel *et al*., 2009). Interestingly, both genes, along with other metal homeostasis actors, are regulated by FIT1 in response to Fe deficiency (Colangelo & Guerinot, 2004). Groups C to E contained many genes typically associated to an Fe deficiency response, some of which were strongly up-regulated upon Zn excess in RR, and to a lower extent in RT. These included genes involved in Fe mobilization through coumarin synthesis (*F6’H1*, *CYP82C4*, *S8H*), uptake (*IRT1*, *FRO2*), trafficking and long distance transport (several *NASs* and *FROs*, *YSL2*, *OPT3*), and the regulation of this response (several *IMAs* and *bHLHs*, *WRKY46*, *MYB10* and *MYB72*) (Kobayashi *et al*., 2019; Hanikenne *et al*., 2021; Riaz & Guerinot, 2021; Vélez-Bermúdez & Schmidt, 2023). In contrast, genes associated with Fe sequestration were solely found among the genes down-regulated by Zn excess, either specifically in RT (*FER2*, *FER3*), in RR (*VTL1*, *2* and *5*) or in both (*FER1*, *FER4*) (Fig. **5**), indicating a need for reduced Fe storage in both root parts (Briat *et al*., 2010; Gollhofer *et al*., 2014).

### Specialized metabolism is up-regulated in RT upon Zn excess

The GO enrichment analysis of the transcriptome data hinted that an increase in specialized metabolite synthesis may take place specifically in RT upon Zn excess (Fig. **4f**). Metabolomic profiling of RT and RR after 48h of Zn excess yielded the detection of 2959 features, of which 215 were unambiguously annotated at family level (Table **S7**) among those ∼12% (RT) and ∼16% (RR) accumulated differentially upon Zn excess. A PCA showed that Zn excess altered the metabolomic profile of both root parts (Fig. **6a**). While most of the metabolomic variation was explained by the root part (PC1, 57% of explained variance), the treatment had a much stronger impact on the metabolome (PC2, 18% of explained variance) than the transcriptome (PC2, 2%, Fig. **4a**).

**Figure 6.**
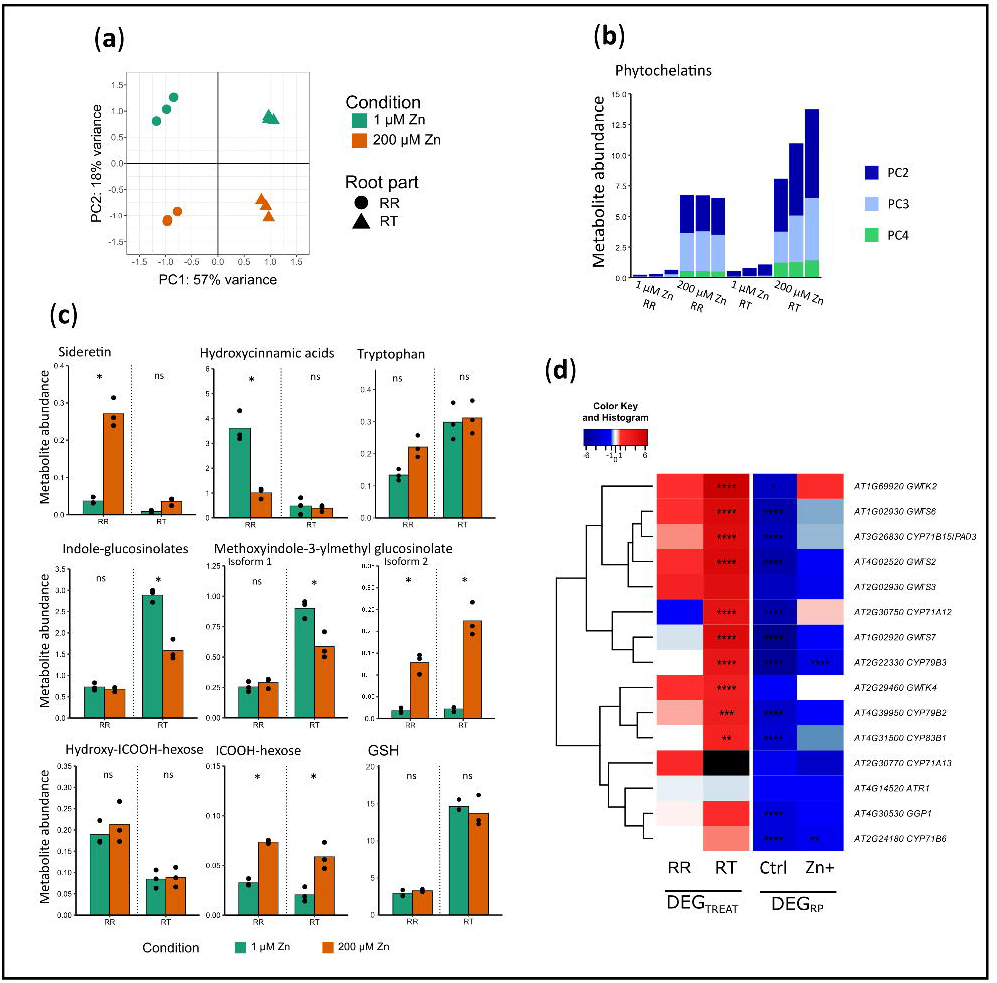
Untargeted metabolomic profiling of the root tip and the remaining root system upon Zn excess in Arabidopsis. Seedlings were germinated and grown for one week in control agar plates (1 µM Zn), then transferred for 48h onto new control (1 µM Zn) or Zn excess (200 µM Zn) plates. Root tip (RT) and remaining root (RR, 2-3 mm segment ∼mid-length of the root) samples were then submitted to untargeted metabolomic. **a.** Principal Component Analysis of the metabolomic data. **b-c.** Phytochelatin (dimer, trimer and tetramer) and other metabolite abundance. **a-c.** Metabolite concentrations were normalized to the total metabolite count of each sample, and results are presented in ‰ of the total metabolite count for each sample. **b.** Each bar represents one replicate. **c.** Black dots are individual replicate values and bars represent means. Stars indicate statistically significant differences between treatments. **d**. Heatmap for the expression of genes of the Trp-derived and camalexin biosynthetic pathway (Fig. **S10**), based on RNA-Seq data. It shows log2(fold change) of gene expression between Zn excess (Zn+) and control conditions (DEG_TREAT_) or between RT and RR (DEG_RP_) after 48h of Zn excess (200 µM Zn). Statistically-significant changes in gene expression (adjusted *p-value* < 0.05) are marked with stars: “ * ” for pval < 0.05, “ ** ” for pval <0.01, “ *** ” for pval < 0.001, “ **** ” for pval < 0.0001.

The abundance of phytochelatins (PCs), known to accumulate in response to Zn excess (Tennstedt *et al*., 2009), increased in RR upon Zn excess (Fig. **6b**). A similar relative increase was observed in RT, indicating that PCs may also participate in Zn chelation in RT. However, the feature annotated as reduced glutathione (GSH), precursor to PCs, remained unchanged by Zn excess (Fig. **6c**).

The abundance of sideretin, a hydroxycoumarin involved in Fe mobilization by roots (Rajniak *et al*., 2018; Clemens, 2019; Paffrath *et al*., 2024), increased in RR but not RT, thereby reflecting the expression profile of its biosynthetic gene (*CYP82C4*) and more generally genes related to Fe-deficiency response and coumarin production (Fig. **6c**, Fig. **5** Group E). The abundance of hydroxycinnamic acids, previously shown to accumulate in Cd-exposed roots of a Cd hypertolerant *Arabidopsis halleri* ecotype (Corso *et al*., 2018), strongly decreased upon Zn excess in RR (Fig. 6c). These compounds, as sideretin, were almost absent from the RT metabolomic profiles, again highlighting differences in metabolomic responses between root parts.

Upon Zn excess, genes encoding enzymes involved in the synthesis of indole-containing compounds and glucosinolates were enriched among genes up-regulated in RT (Fig. **4f**). While the abundance of tryptophan (Trp), a precursor for the biosynthesis of these compounds (Fig. **S10**), remained unchanged upon Zn excess in both RT and RR, total indole-glucosinolates decreased in RT. However, an unknown isoform of methoxyindole-3-ylmethyl glucosinolate was more abundant in both RT and RR upon Zn excess (Isoform 2 in Fig. **6c**), whereas the abundance ofindole-3-carboxylic acid (ICOOH)-hexose was increased in both root parts (Fig. **6c**). The increased levels of ICOOH-hexose and the second isoform of methoxyindole-3-ylmethyl glucosinolate, endpoints of biosynthetic pathways from Trp, reflected the transcriptomic changes of several of their biosynthesis genes (Fig. **6d**, Fig. **S10**).

### Camalexin synthesis is increased upon Zn excess

Camalexin, a specialized metabolite derived from Trp and known for its role in defense against fungal pathogen attack (Mucha *et al*., 2019), was highlighted in the GO enrichment analysis as being specifically enriched in the RT in response to Zn excess (Fig. **4f**). Several genes involved in camalexin biosynthesis, including *PAD3*, which encodes the final enzyme in this pathway (Fig. **S10**), were induced by Zn excess in RT (Fig. **6d**, Fig. **7a**). Although camalexin was not annotated in the untargeted metabolomics dataset, targeted LC-MS analysis revealed strongly increased camalexin concentrations in both Arabidopsis roots and shoots upon Zn excess (Fig. **7b**).

**Figure 7.**
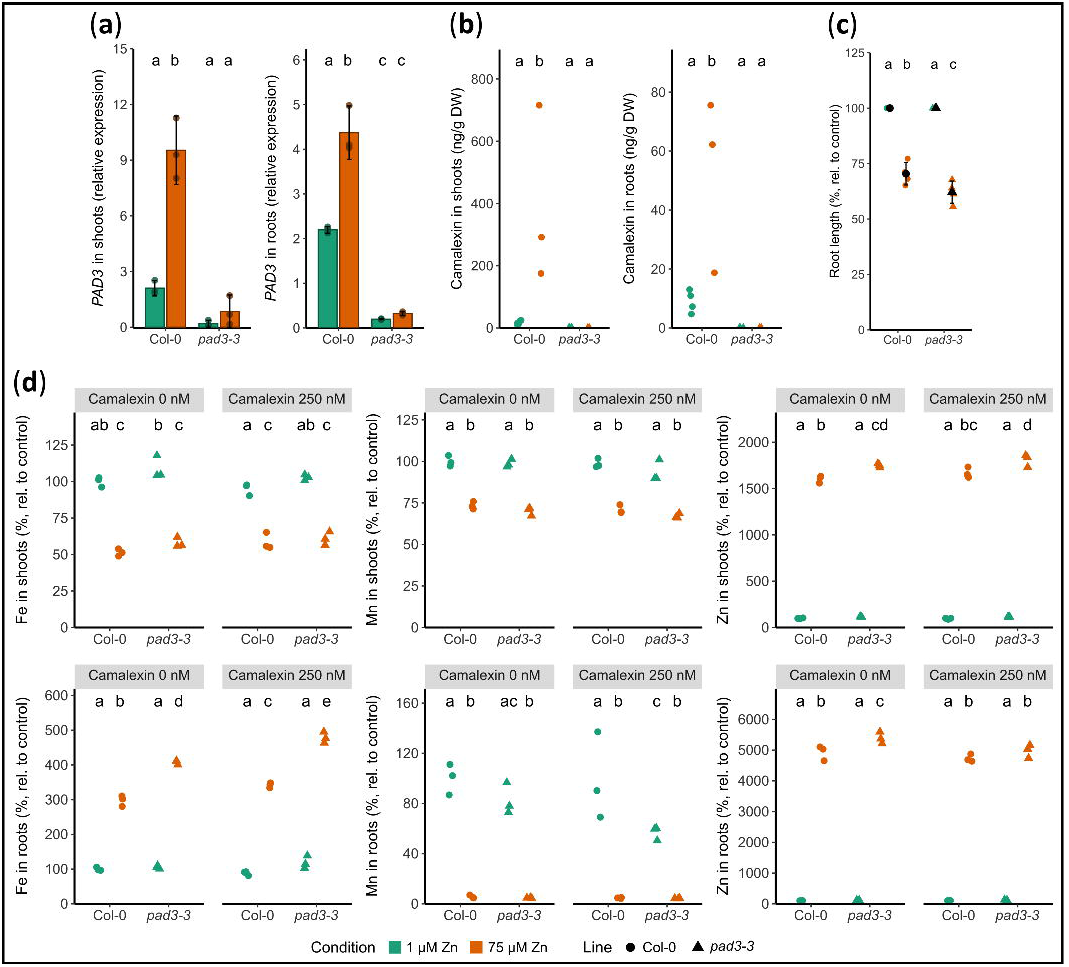
Responses of a *pad3* mutant to Zn excess on Arabidopsis. Col-0 and *pad3-3* knocked-out mutant seedlings were germinated and grown for 11 days on agar plates with either control conditions (1 µM Zn) or Zn excess (75 µM Zn) (a-d), and control (0 nM) or exogenous camalexin treatment (250 nM) (d). a. *PAD3* expression assed by RT-qPCR, from 3 biological replicates. b. Camalexin quantification in roots and shoots of the two genotypes from 4 biological replicates. c. Primary root growth after 11 days of growth, relative to their respective control condition, from 4 independent replicates. d. Elemental concentrations relative to their concentrations in Col-0 in control condition, from 3 biological replicates. Individual replicate values or shapes (a-d) and whiskers representing mean values and standard deviations (a) are presented. Different letters correspond to significantly different groups (ANOVA type I, with Tukey test correction, *p-value* < 0.05). Root length from the other *pad3* mutants are presented in Fig. S11, for other element concentrations in Fig. S12.

To examine the contribution of camalexin to Zn homeostasis, a T-DNA insertion mutant (*pad3-3*) was studied. The mutant displayed strongly reduced *PAD3* expression (Fig. **7a**) and no camalexin synthesis (Fig. **7b**) neither in roots nor shoots, regardless of Zn conditions. Primary root growth inhibition was more pronounced in *pad3-3* than in Col-0 seedlings 24h and 48h after transfer on 200 µM Zn excess (Fig. **S11**). Moreover, when germinated and grown directly on 75 µM Zn medium, a greater reduction in primary root growth was observed in *pad3-3* compared to Col-0 (Fig. **7c**). This reduced growth phenotype upon Zn excess was confirmed in two additional *pad3* mutant alleles (Fig. **S11**).

In control conditions, no significant differences were observed in the ionome profile of the *pad3-3* mutant compared to Col-0, except for an increase of Mg in shoots. Conversely, upon 75 µM Zn excess, the *pad3-3* mutant displayed increased concentrations of Fe and Zn in roots and Fe in shoots (Fig. **7d**, Fig. **S12**). Next, the effect of camalexin on these ionome phenotypes was examined. Adding camalexin (250 nM) to the control medium (1 µM Zn) had limited effect on the ionome of both Col-0 and *pad3-3*, except for a significant reduction in Mn concentration in *pad3-3* roots. However, upon Zn excess (75 µM), camalexin-treated Col-0 and *pad3-3* seedlings displayed increased root Fe concentration compared to untreated seedlings. Furthermore, mutant seedlings exposed to both Zn excess and camalexin showed a significant decrease in root Zn concentration, down to Col-0 levels (Fig. **7d**).

Finally, the expression of a panel of marker genes was examined in the mutant to further characterize its defects Zn and Fe homeostasis. Among those, the *HMA2* and *FRD3* genes, involved in Zn and Fe translocation to shoots, respectively, were significantly less expressed in roots of the mutant compared to Col-0 in control conditions (Fig. **S13**).

## Discussion

Differentiated root tissues display specific traits, such as large vacuoles and apoplastic barriers, which are absent in meristematic cells and could significantly affect nutrient homeostasis. Here, we used a multi-omics approach to comprehensively examine the specific responses of the RT to Zn excess, in contrast to RR. This revealed distinct Zn distribution, uptake, storage capacities and Fe homeostasis in the RT. We further identified a strong and specific metabolic response to Zn in the RT that contributes to Zn tolerance. Together with a report on the impact of Cd in a similar experimental design (Richtmann *et al*.), our data highlight both similarities and differences in the response to excess of essential (Zn) and non-essential (Cd) metals.

### Zn uptake, storage and chelation capacities differ in RT and RR

Our study shows that Zn homeostasis processes are partitioned between RT and RR, with a uniform Zn distribution across the different RAM cell layers, and a prevalence in the vascular cylinder of differentiated roots (Fig. **3**) (Sinclair *et al*., 2007; Persson *et al*., 2016; Giehl *et al*., 2023). Upon Zn excess, Zn concentrations strongly increased in both RR and RT, but less so in the RT, where it only was ∼64% of that in the RR (Fig. **2**), a pattern similar to the reduced Cd accumulation observed in the RT (Richtmann *et al*.). These distinct distribution and accumulation in the RT correlated with a globally lower expression of known Zn homeostasis genes (Fig. **5**), suggesting reduced Zn uptake, vacuolar storage and chelation capacities.

Indeed, several Zn-responsive *ZIP* genes (*ZIP3*, *ZIP5* and *IRT3*, Fig. **5** Group B), thought to be responsible for primary Zn entry in root cells (Sinclair & Krämer, 2012; Lee *et al*., 2021), were only marginally expressed in RT under both control and Zn excess conditions, whereas Zn excess typically reduced their expression in RR (Talke *et al*., 2006; Lin *et al*., 2009). Despite this lower uptake capacity in RT, Zn concentrations in RT and RR were similar in control conditions (Fig. **2**), suggesting the existence of a Zn transfer mechanism from RR to RT. Such a hypothesis implies that upon Zn excess, this yet unknown transfer mechanism is perturbed, further contributing to lower Zn accumulation in RT.

Moreover, several observations are consistent with limited capacity for Zn vacuolar storage and chelation in RT, as this compartment only represents ∼20% of the volume in RAM cells (Cui *et al*., 2019) compared to ∼90% in mature cells (Kaiser & Scheuring, 2020). This is further supported by the lower expressions of the 4 *NAS* genes, involved in nicotianamine (NA) synthesis (Clemens, 2019) and *ZIF1*, encoding a vacuolar NA transporter (Haydon *et al*., 2012), in RT compared to RR (Fig. **5**, Table **S2**). *NAS4* induction by Zn excess was further limited to RR. The *MTP1*, *MTP3* and *HMA3* genes, encoding vacuolar Zn transporters (Desbrosses-Fonrouge *et al*., 2005; Arrivault *et al*., 2006; Morel *et al*., 2009), were either similarly or less expressed in RT compared to RR, respectively, with *MTP3* and *HMA3* induction by high Zn (Colangelo & Guerinot, 2004; Arrivault *et al*., 2006) being again limited to RR (Fig. **5**). The >7-fold increase in Zn chelator phytochelatin accumulation upon Zn excess (Fig. **6b**) indicated a high cytosolic Zn concentration in the RT (Tennstedt *et al*., 2009).

### Distinct Fe homeostasis actors respond to Zn excess in RT and RR

At the transcriptional level, Zn excess had a comparatively larger impact on Fe than on Zn homeostasis genes, leading to a strong Fe deficiency response in RR (Fig. **4e-f**, Fig. **5** Groups C-E), particularly affecting genes downstream of the FIT transcription factor (Schwarz and Bauer, 2020). Such a response is typical in Arabidopsis, stemming from Zn and Fe homeostasis interactions through shared metal transporters and chelators (Fukao *et al*., 2011; Shanmugam *et al*., 2011; Hanikenne *et al*., 2021). The increased synthesis of sideretin, a coumarin compound that aids in Fe solubilization and uptake, reflected this response in the RR metabolome (Murgia *et al*., 2011; Clemens & Weber, 2016; Tsai *et al*., 2018; Paffrath *et al*., 2024) (Fig. **6c**). This response likely maintained Fe accumulation in RR during short-term (24-48h) Zn excess (Fig. **2b**), and contrasts with reduced sideretin accumulation observed upon Cd exposure (Richtmann *et al*.).

In control conditions, Fe homeostasis genes were generally less expressed and displayed a more limited Fe deficiency response to Zn excess in RT than in RR. Up-regulation occurred only for a transcriptional regulator (*bHLH39*), an Fe phloem transporter (*OPT3*), phloem-localized IRONMAN signaling peptides (*IMAs*) (Riaz & Guerinot, 2021; Vélez-Bermúdez & Schmidt, 2023), and *IRT1* and *IRT2* (Vert *et al*., 2009; Spielmann & Vert, 2021). Furthermore, *FER2* and *FER3* genes, encoding ferritins for Fe storage, were specifically down-regulated in the RT upon Zn excess (Fig. **5**). Together with lower Zn accumulation in RT, this appeared sufficient to maintain Fe concentration in RT upon Zn excess (Fig. **2**). In addition to the aforementioned reduced expression of the Zn-responsive *ZIP* genes, the lower expression of *IRT1* in RT may further limit Zn accumulation. Despite a complex posttranslational regulation by non-Fe metals (*e.g.* Zn) (Spielmann and Vert, 2021), IRT1, the main Fe uptake transporter in Arabidopsis, is believed to substantially contribute to Zn uptake by roots (Shanmugam *et al*., 2011; Hanikenne *et al*., 2021; Merlot *et al*., 2021; Spielmann *et al*., 2024).

RT and RR further differed in their distinct Fe distribution patterns. In RR, Fe mainly accumulated in the epidermis, as described previously in older roots (Persson *et al*., 2016). In RAM, Fe accumulated mainly in the future cortical ring cells (Fig. **3**), consistent with previous observations (Reyt *et al*., 2015). Contrary to most Zn and Fe homeostasis genes, some Fe homeostasis genes had similar expression levels in RT and RR (Group A, Fig. **5**, including several *YSLs* and *NRAMPs*). Whether they determine this cortical ring distribution of Fe in the RAM will be an interesting question to pursue. Conversely, the RT-specific *FER2-3* genes likely have a negligible role in Fe distribution between RAM cell types, as an Fe cortical ring was also observed in a *fer1fer3fer4* Arabidopsis triple mutant (Reyt *et al*., 2015).

### Root growth processes are impacted by Zn excess

In Arabidopsis, imbalance in metal supply (either excess or deficiency) impairs primary root growth (Gruber *et al*., 2013; Lešková *et al*., 2017). Such growth reduction is thought to reflect altered resource allocation upon stress (Bechtold & Field, 2018; Carrera *et al*., 2018; van Dijk *et al*., 2022). Root growth processes, consisting of meristematic divisions, cell elongation and finally differentiation, were all impacted by Zn excess: the RAM and elongation zone were shortened, with differentiation initiated closer to the root tip (Fig. **1**). The reduced number of cells in the RAM suggested a negative impact of Zn on cell division. Although cell cycle genes were not among the differentially expressed genes (DEGs) in the RT upon Zn excess (Fig. **S8**), this does not contradict this interpretation, as cell cycle regulators are mostly modulated post-transcriptionally (Inzé & De Veylder, 2006; Genschik *et al*., 2014; Yang *et al*., 2017). RAM and elongation zone delineations are controlled by distinct hormonal and redox gradients (Mase & Tsukagoshi, 2021; Zluhan-Martínez *et al*., 2021) [e. g. by opposite auxin-cytokinin gradients, with auxin levels peaking in QC and decreasing upwards in the root (as reviewed in Schaller *et al*., 2015)]. Consistent with previous findings that Zn excess alters auxin signaling in the RAM (Zhang *et al*., 2018; Wang *et al*., 2021), we observed a reduced RAM size (Fig. **1c-e**). At the transcriptional level, auxin response in the RT was limited, except for the differential expression of several small auxin up-regulated RNA (*SAUR* and *SAUR-like*) upon Zn excess (Table **S2**, Table **S3**). *SAUR* genes contribute to the control of cell elongation, often in response to environmental cues (Stortenbeker & Bemer, 2019).

Despite the major morphological impact of Zn excess on roots, the root part was the main determinant of gene differential expression, reflecting a partitioning of developmental processes and functions between RT and RR (Fig. **4a**) (discussed in Richtmann *et al*.). Moreover, ∼30% of the DEGs in the RT upon Zn excess were associated with root differentiation (Table **S5**), further highlighting a Zn-triggered shift in the positioning of the division, elongation and differentiation zones in the RT (Fig. **1g**).

### A specific metabolomic response takes place in the RT upon Zn excess

Strikingly, Zn excess impacted on the root metabolome (Fig. **6a**) much more than the transcriptome (Fig. **4a**), 18% vs 2% of explained variance in PCA, respectively, and this response was quite divergent from Cd stress, particularly in the RT (Fig. **S14**) (Richtmann *et al*.). Indeed, in the RT, 44% and 58% of the features differentially accumulated under Zn and Cd treatments, respectively, were unique to each condition, compared to 27% and 39% in the RR. Notably, a large Trp-derived specialized metabolic pathway, including indole glucosinolates, indole carboxylic acid derivatives, camalexin and indole-3-carbonylnitrile, was massively up-regulated specifically in the RT upon Zn excess, (Fig. **4f**, Fig. **6d**, Fig. **S10**), but not upon Cd stress (Richtmann *et al*.). This specialized pathway is controlled by the WRK33 transcription factor (Petersen *et al*., 2008; Barco *et al*., 2019; Cabot *et al*., 2019), whose transcript was also upregulated in RT upon Zn excess.

These metabolites share the intermediate indole-3-acetaldoxime (IAOx, Fig. **S10**), whose production from Trp is also promoted by Zn exposure in peas (Horák *et al*., 1976). The common requirement for IAOx, but also GSH and possibly NADPH, leads to substrate competition among the branches of the pathway (Malka & Cheng, 2017; Vik *et al*., 2018; Pastorczyk *et al*., 2020). Increased PC synthesis upon Zn excess (Fig. **6b**) may further divert GSH away from these pathways. This could explain why the upregulation of several genes in these biosynthetic pathways (Fig. **6d**) resulted in either increased (*e.g.* ICOOH-hexose, camalexin) or decreased (*e.g.* methoxyindol-3-ylmethylglucosinolate) derived metabolite accumulation upon Zn excess (Fig. **6c**, Fig. **7b**).

Although Zn is not a redox-active ion, oxidative stress-like responses were observed in Arabidopsis roots exposed to Zn excess (Remans *et al*., 2012; De Smet *et al*., 2015; Cuypers *et al*., 2016), which could impact the redox balance in the RT, crucial for cell division and RAM size determination (Tsukagoshi *et al*., 2010; Mase & Tsukagoshi, 2021). RT-specific metabolic changes may contribute to maintain its redox status. The specific down-regulation of *FER2* and *FER3* in the RT, encoding ferritins crucial for protecting cells from Fe-triggered oxidative stress (Ravet *et al*., 2009), could further help to maintain an adequate Fe/Zn ratio in RAM cells, essential for redox balance (Shanmugam *et al*., 2011; Reyt *et al*., 2015; Scheepers *et al*., 2020).

### Increased camalexin synthesis contributes to the Zn excess response

Camalexin, a major phytoalexin in Brassicaceae, is massively synthesized and secreted as a defense response to pathogen attacks (Tsuji *et al*., 1992; Khare *et al*., 2017; Tewes *et al*., 2018). Although its molecular mode of action remains to be established, its antifungal properties are believed to stem from its antioxidant capacities as, *in vitro*, camalexin scavenges radicals and even displays ferric reducing antioxidant power similar to ascorbic acid (Manasa & Chitra, 2020).

So far, Zn and camalexin have been used as a model system to test trade-off or joint effect hypotheses of metal and chemical defense against pathogens in leaves, with different outcomes (Martos *et al*., 2016; Tewes *et al*., 2018; Cabot *et al*., 2019). In the Zn and Cd hyperaccumulator *A. halleri* (Merlot *et al*., 2021), a trade-off was observed: high Zn and Cd reduced shoot camalexin concentration without impacting pathogen resistance (Tewes *et al*., 2018). Conversely, high Zn increased pathogen resistance in Arabidopsis, which was not conserved in a *pad3* mutant, supporting a joint effect hypothesis (Martos *et al*., 2016). Here, we showed that Zn tolerance, along with Fe and Mn homeostasis, were affected in the camalexin-free *pad3-3* Arabidopsis mutant (Fig. **7d**, Fig. **S12**), suggesting a role for camalexin synthesis in metal homeostasis. Increased camalexin synthesis appears to be specific to Zn excess response, as *PAD3* expression was not induced under Cd excess (Richtmann *et al*.), or Fe deficiency (Fig. **S15**).

At the molecular level, the interplay between Zn and camalexin may result from the impact of Zn on multiple intricate processes. The Zn-induced upregulation of *PAD3* was suggested to be caused by a stimulation of the jasmonic acid and ethylene signaling pathways (Martos *et al*., 2016), which we also observed in the RT (Table **S3**). Additionally, ROS production and altered GSH status, common features of the plant response to biotic and abiotic stresses (Castro *et al*., 2021; van Dijk *et al*., 2022), are also major triggers for increased camalexin synthesis (Glawischnig, 2007). The sensitivity of the *pad3-3* mutant to Fe excess, a condition known to trigger oxidative stress (Reyt *et al*., 2015; Li *et al*., 2019), further supports this hypothesis (Fig. **S16**). In addition to a direct antioxidant activity, camalexin may also bind Zn (and other metals), similarly to the proposed role of flavonoids in Cd tolerance (Corso *et al*., 2018; Richtmann *et al*.), but so far we have not been able to collect experimental evidence for this. A working hypothesis is that a limited capacity to store excess Zn in the small vacuoles of the RT cells (Fig. **5**) is compensated for by camalexin, possibly acting as an antioxidant, protecting the RT and regulating the redox status required for cell division in the RAM. As such, this hypothesis suggests that camalexin plays a crucial role in the RT, beyond its well-described function in foliar pathogen protection. While camalexin, together with other Trp-derived metabolites, is secreted *via* PEN3 and PDR12 transporters upon pathogen infection (He *et al*., 2019), the absence of regulation of *PEN3* and *PDR12* in RT upon Zn excess (Table **S3**), suggests that camalexin was not excreted in the roots upon Zn excess but rather acted intracellularly.

In conclusion, this study describes the distinct metal homeostasis mechanisms taking place in RT and RR in response to Zn excess, highlighting distinct partitioning of Zn and Fe homeostasis. It emphasizes how a Trp-derived specialized metabolic response in the RT may contribute to the Zn excess response by acting on redox homeostasis to protect cell division, given the overall limited metal transport and storage capacities in the RT. Hence, future research should aim to dissect how these Trp-derived defense metabolites contribute to root Zn tolerance, and link Zn homeostasis to redox control and biotic defense pathways (Zamioudis *et al*., 2015; Martos *et al*., 2016; Tewes *et al*., 2018; Scheepers *et al*., 2020). Such crosstalk is reminiscent of the function of defensin(-like) proteins (Kimura *et al*., 2023; Nguyen *et al*., 2023).

## Supporting information

Thiebaut_Supporting information

## Acknowledgements

We thank A. Degueldre and B. Bosman (ULiège, Belgium) for technical support in ICP-OES analyses, L. Karim and Dr. W. Coppieters (ULiège, Belgium) for performing the RNA sequencing, as well as Dr. M. Corso (Institut Jean-Pierre Bourgin, France) for support in metabolomics data analyses. We thank Dr. A. Assunção (UCopenhagen, Denmark) for support in plant culture and discussions, as well as Dr. S. Fanara (ULiège, Belgium) who provided the RT-qPCR Fe deficiency samples. We equally thank Prof. L. Veylder (UGent, Belgium) for helpful discussions at the start of the project. Funding was provided by the “Fonds de la Recherche Scientifique-FNRS” (PDR-T0120.18, PDR-T.0104.22 to M.H. and N.V.), the COST ACTION 19116 PLANTMETALS, and the University of Bayreuth (to SC). The IJPB benefits from the support of Saclay Plant Sciences-SPS (ANR-17-EUR-0007). This work has benefited from the support of IJPB’s Plant Observatory platform PO-Chem.

## Author contributions

MH, NV, and NT and SC designed the research. NT, LRi, MSa, DPP, AR, MSc, SB and LRe performed experiments. NT, LRi, MSa, DPP, SB, LRe, SC, MH and NV analyzed the data. NT, MSa and LRi made the figures. NT, MH, MSa and NV wrote the manuscript. All authors read and approved the manuscript.

## Data availability

The RNA-Seq reads have been deposited in the National Center for Biotechnology Information (NCBI) Sequence Read Archive (SRA) Database with BioProject identification number (submission in progress). The metabolomic data and metadata have been deposited at the MassiVE data repository portal with the identifier (submission in progress). The other data that support the findings of this study are available from the corresponding authors upon reasonable request.

## Literature

Ahuja I, Kissen R, Bones AM. 2012. Phytoalexins in defense against pathogens. Trends in Plant Science 17: 73–90.

Amini S, Arsova B, Hanikenne M. 2022. The molecular basis of zinc homeostasis in cereals. Plant Cell and Environment 45: 1339–1361.

Arrivault S, Senger T, Krämer U. 2006. The Arabidopsis metal tolerance protein AtMTP3 maintains metal homeostasis by mediating Zn exclusion from the shoot under Fe deficiency and Zn oversupply. Plant Journal 46: 861–879.

Assunção AGL, Herrero E, Lin YF, Huettel B, Talukdar S, Smaczniak C, Immink RGH, Van Eldik M, Fiers M, Schat H, et al. 2010. Arabidopsis thaliana transcription factors bZIP19 and bZIP23 regulate the adaptation to zinc deficiency. Proceedings of the National Academy of Sciences of the United States of America 107: 10296–10301.

Barberon M. 2017. The endodermis as a checkpoint for nutrients. New Phytologist 213: 1604–1610.

Barberon M, Vermeer JEM, De Bellis D, Wang P, Naseer S, Andersen TG, Humbel BM, Nawrath C, Takano J, Salt DE, et al. 2016. Adaptation of Root Function by Nutrient-Induced Plasticity of Endodermal Differentiation. Cell 164: 447–459.

Barco B, Kim Y, Clay NK. 2019. Expansion of a core regulon by transposable elements promotes Arabidopsis chemical diversity and pathogen defense. Nature Communications 10: 1–12.

Bechtold U, Field B. 2018. Molecular mechanisms controlling plant growth during abiotic stress. Journal of Experimental Botany 69: 2753–2758.

Boutet S, Barreda L, Perreau F, Totozafy JC, Mauve C, Gakière B, Delannoy E, Martin-Magniette ML, Monti A, Lepiniec L, et al. 2022. Untargeted metabolomic analyses reveal the diversity and plasticity of the specialized metabolome in seeds of different Camelina sativa genotypes. Plant Journal 110: 147– 165.

Briat JF, Duc C, Ravet K, Gaymard F. 2010. Ferritins and iron storage in plants. Biochimica et Biophysica Acta - General Subjects 1800: 806–814.

Bruno L, Pacenza M, Forgione I, Lamerton LR, Greco M, Chiappetta A, Bitonti MB. 2017. In Arabidopsis thaliana cadmium impact on the growth of primary root by altering SCR expression and auxin-cytokinin cross-talk. Frontiers in Plant Science 8: 1–13.

Cabot C, Martos S, Llugany M, Gallego B, Tolrà R, Poschenrieder C. 2019. A Role for Zinc in Plant Defense Against Pathogens and Herbivores. Frontiers in Plant Science 10: 1–15.

Cai G, Ahmed MA. 2022. The role of root hairs in water uptake: Recent advances and future perspectives. Journal of Experimental Botany 73: 3330–3338.

Carrera D, Oddsson S, Grossmann J, Trachsel C, Streb S. 2018. Comparative Proteomic Analysis of Plant Acclimation to Six Different Long-Term Environmental Changes. Plant and Cell Physiology 59: 510–526.

Castaings L, Caquot A, Loubet S, Curie C. 2016. The high-affinity metal Transporters NRAMP1 and IRT1 Team up to Take up Iron under Sufficient Metal Provision. Scientific Reports 6: 1–11.

Castro B, Citterico M, Kimura S, Stevens DM, Wrzaczek M, Coaker G. 2021. Stress-induced reactive oxygen species compartmentalization, perception and signalling. Nature Plants 2021 7:4 7: 403–412.

Clemens S. 2019. Metal ligands in micronutrient acquisition and homeostasis. Plant Cell and Environment 42: 2902–2912.

Clemens S. 2022. The cell biology of zinc. Journal of Experimental Botany 73: 1688–1698.

Clemens S, Deinlein U, Ahmadi H, Höreth S, Uraguchi S. 2013. Nicotianamine is a major player in plant Zn homeostasis. BioMetals 26: 623–632.

Clemens S, Weber M. 2016. The essential role of coumarin secretion for Fe acquisition from alkaline soil. Plant Signaling and Behavior 11: 1–6.

Colangelo EP, Guerinot M Lou. 2004. The essential basic helix-loop-helix protein FIT1 is required for the iron deficiency response. Plant Cell 16: 3400–3412.

Corso M, Schvartzman MS, Guzzo F, Souard F, Malkowski E, Hanikenne M, Verbruggen N. 2018. Contrasting cadmium resistance strategies in two metallicolous populations of Arabidopsis halleri. New Phytologist 218: 283–297.

Cui Y, Cao W, He Y, Zhao Q, Wakazaki M, Zhuang X, Gao J, Zeng Y, Gao C, Ding Y, et al. 2019. A whole-cell electron tomography model of vacuole biogenesis in Arabidopsis root cells. Nature Plants 5: 95– 105.

Cuypers A, Hendrix S, dos Reis RA, De Smet S, Deckers J, Gielen H, Jozefczak M, Loix C, Vercampt H, Vangronsveld J, et al. 2016. Hydrogen peroxide, signaling in disguise during metal phytotoxicity. Frontiers in Plant Science 7: 470.

Desbrosses-Fonrouge AG, Voigt K, Schröder A, Arrivault S, Thomine S, Krämer U. 2005. Arabidopsis thaliana MTP1 is a Zn transporter in the vacuolar membrane which mediates Zn detoxification and drives leaf Zn accumulation. FEBS Letters 579: 4165–4174.

van Dijk JR, Kranchev M, Blust R, Cuypers A, Vissenberg K. 2022. Arabidopsis root growth and development under metal exposure presented in an adverse outcome pathway framework. Plant Cell and Environment 45: 737–750.

Dubeaux G, Neveu J, Zelazny E, Vert G. 2018. Metal Sensing by the IRT1 Transporter-Receptor Orchestrates Its Own Degradation and Plant Metal Nutrition. Molecular Cell 69: 953–964.e5.

Fukao Y, Ferjani A, Tomioka R, Nagasaki N, Kurata R, Nishimori Y, Fujiwara M, Maeshima M. 2011. iTRAQ analysis reveals mechanisms of growth defects due to excess zinc in Arabidopsis. Plant Physiology 155: 1893–1907.

Genschik P, Marrocco K, Bach L, Noir S, Criqui MC. 2014. Selective protein degradation: A rheostat to modulate cell-cycle phase transitions. Journal of Experimental Botany 65: 2603–2615.

Giehl RFH, Flis P, Fuchs J, Gao Y, Salt DE, von Wirén N. 2023. Cell type-specific mapping of ion distribution in Arabidopsis thaliana roots. Nature Communications 14: 1–12.

Glawischnig E. 2007. Camalexin. Phytochemistry 68: 401–406.

Gollhofer J, Timofeev R, Lan P, Schmidt W, Buckhout TJ. 2014. Vacuolar-iron-transporter1-like proteins mediate iron homeostasis in Arabidopsis. PLoS ONE 9: e110468.

Gruber BD, Giehl RFH, Friedel S, von Wirén N. 2013. Plasticity of the Arabidopsis root system under nutrient deficiencies. Plant Physiology 163: 161–179.

Hanikenne M, Esteves SM, Fanara S, Rouached H. 2021. Coordinated homeostasis of essential mineral nutrients: A focus on iron. Journal of Experimental Botany 72: 2136–2153.

Haydon MJ, Kawachi M, Wirtz M, Hillmer S, Hell R, Krämer U. 2012. Vacuolar nicotianamine has critical and distinct roles under iron deficiency and for zinc sequestration in Arabidopsis. Plant Cell 24: 724–737.

He Y, Xu J, Wang X, He X, Wang Y, Zhou J, Zhang S, Meng X. 2019. The Arabidopsis Pleiotropic Drug Resistance Transporters PEN3 and PDR12 Mediate Camalexin Secretion for Resistance to Botrytis cinerea. The Plant cell 31: 2206–2222.

Henriques R, Jásik J, Klein M, Martinoia E, Feller U, Schell J, Pais MS, Koncz C. 2002. Knock-out of Arabidopsis metal transporter gene IRT1 results in iron deficiency accompanied by cell differentiation defects. Plant Molecular Biology 50: 587–597.

Horák V, Tròka I, Stefl M. 1976. The Influence of Zn 2+ Ions on the Tryptophan Biosynthesis in Plants V. Biologia Plantarum 18: 393–396.

Inzé D, De Veylder L. 2006. Cell Cycle Regulation in Plant Development. Annual Review of Genetics 40: 77–105.

Kaiser S, Scheuring D. 2020. To Lead or to Follow: Contribution of the Plant Vacuole to Cell Growth. Frontiers in Plant Science 11: 8–13.

Kaur H, Garg N. 2021. Zinc toxicity in plants: a review. Planta 253: 1–28.

Khare D, Choi H, Huh SU, Bassin B, Kim J, Martinoia E, Sohn KH, Paek KH, Lee Y, Chrispeels MJ. 2017. Arabidopsis ABCG34 contributes to defense against necrotrophic pathogens by mediating the secretion of camalexin. Proceedings of the National Academy of Sciences of the United States of America 114: E5712–E5720.

Kimura S, Vaattovaara A, Ohshita T, Yokoyama K, Yoshida K, Hui A, Kaya H, Ozawa A, Kobayashi M, Mori IC, et al. 2023. Zinc deficiency-induced defensin-like proteins are involved in the inhibition of root growth in Arabidopsis. The Plant Journal 115: 1071–1083.

Kobayashi T, Nozoye T, Nishizawa NK. 2019. Iron transport and its regulation in plants. Free Radical Biology and Medicine 133: 11–20.

Kondo Y, Tamaki T, Fukuda H. 2014. Regulation of xylem cell fate. Frontiers in Plant Science 5: 1–6.

Lee S, Lee J, Ricachenevsky FK, Punshon T, Tappero R, Salt DE, Guerinot M Lou. 2021. Redundant roles of four ZIP family members in zinc homeostasis and seed development in Arabidopsis thaliana. Plant Journal: 1162–1173.

Leitner D, Klepsch S, Ptashnyk M, Marchant A, Kirk GJD, Schnepf A, Roose T. 2010. A dynamic model of nutrient uptake by root hairs. New Phytologist 185: 792–802.

Leonardo B, Emanuela T, Letizia MM, Antonella M, Marco M, Fabrizio A, Beatrice BM, Adriana C. 2021. Cadmium affects cell niches maintenance in *Arabidopsis thaliana* post-embryonic shoot and root apical meristem by altering the expression of WUS/WOX homolog genes and cytokinin accumulation. Plant Physiology and Biochemistry 167: 785–794.

Lešková A, Giehl RFH, Hartmann A, Fargašová A, von Wirén N. 2017. Heavy Metals Induce Iron Deficiency Responses at Different Hierarchic and Regulatory Levels. Plant Physiology 174: 1648–1668.

Li B, Sun L, Huang J, Göschl C, Shi W, Chory J, Busch W. 2019. GSNOR provides plant tolerance to iron toxicity via preventing iron-dependent nitrosative and oxidative cytotoxicity. Nature Communications 2019 10:1 10: 1–13.

Lilay GH, Thiébaut N, du Mee D, Assunção AGL, Schjoerring JK, Husted S, Persson DP. 2024. Linking the key physiological functions of essential micronutrients to their deficiency symptoms in plants. New Phytologist 242: 881–902.

Lin YF, Liang HM, Yang SY, Boch A, Clemens S, Chen CC, Wu JF, Huang JL, Yeh KC. 2009. Arabidopsis IRT3 is a zinc-regulated and plasma membrane localized zinc/iron transporter. New Phytologist 182: 392–404.

Malka SK, Cheng Y. 2017. Possible interactions between the biosynthetic pathways of indole glucosinolate and Auxin. Frontiers in Plant Science 8: 1–14.

Manasa K, Chitra V. 2020. Evaluation of in-vitro antioxidant activity of camalexin-a novel anti-parkinson’s agent. Research Journal of Pharmacy and Technology 13: 578–582.

Martos S, Gallego B, Cabot C, Llugany M, Barceló J, Poschenrieder C. 2016. Zinc triggers signaling mechanisms and defense responses promoting resistance to Alternaria brassicicola in Arabidopsis thaliana. Plant Science 249: 13–24.

Mase K, Tsukagoshi H. 2021. Reactive Oxygen Species Link Gene Regulatory Networks During Arabidopsis Root Development. Frontiers in Plant Science 12: 1–15.

Merlot S, Sanchez Garcia de LaTorre V, Hanikenne M. 2021. Physiology and Molecular Biology of Trace Element Hyperaccumulation. In: Agromining: Farming for Metals. 155–182.

Morel M, Crouzet J, Gravot A, Auroy P, Leonhardt N, Vavasseur A, Richaud P. 2009. AtHMA3, a P1B-ATPase allowing Cd/Zn/Co/Pb vacuolar storage in Arabidopsis. Plant Physiology 149: 894–904.

Van De Mortel JE, Villanueva LA, Schat H, Kwekkeboom J, Coughlan S, Moerland PD, Van Themaat EVL, Koornneef M, Aarts MGM. 2006. Large expression differences in genes for iron and zinc homeostasis, stress response, and lignin biosynthesis distinguish roots of Arabidopsis thaliana and the related metal hyperaccumulator Thlaspi caerulescens. Plant Physiology 142: 1127–1147.

Mucha S, Heinzlmeir S, Kriechbaumer V, Strickland B, Kirchhelle C, Choudhary M, Kowalski N, Eichmann R, Hückelhoven R, Grill E, et al. 2019. The Formation of a Camalexin Biosynthetic Metabolon. The Plant cell 31: 2697–2710.

Müller J, Toev T, Heisters M, Teller J, Moore KL, Hause G, Dinesh DC, Bürstenbinder K, Abel S. 2015. Iron-Dependent Callose Deposition Adjusts Root Meristem Maintenance to Phosphate Availability. Developmental Cell 33: 216–230.

Murgia I, Tarantino D, Soave C, Morandini P. 2011. Arabidopsis CYP82C4 expression is dependent on Fe availability and circadian rhythm, and correlates with genes involved in the early Fe deficiency response. Journal of Plant Physiology 168: 894–902.

Nguyen NN, Lamotte O, Alsulaiman M, Ruffel S, Krouk G, Berger N, Demolombe V, Nespoulous C, Dang TMN, Aimé S, et al. 2023. Reduction in PLANT DEFENSIN 1 expression in Arabidopsis thaliana results in increased resistance to pathogens and zinc toxicity. Journal of Experimental Botany 74: 5374– 5393.

Paffrath V, Tandron Moya YA, Weber G, von Wirén N, Giehl RFH. 2024. A major role of coumarin-dependent ferric iron reduction in strategy I-type iron acquisition in Arabidopsis. The Plant Cell 36: 642–664.

Pastorczyk M, Kosaka A, Piślewska-Bednarek M, López G, Frerigmann H, Kułak K, Glawischnig E, Molina A, Takano Y, Bednarek P. 2020. The role of CYP71A12 monooxygenase in pathogen-triggered tryptophan metabolism and Arabidopsis immunity. New Phytologist 225: 400–412.

Perilli S, Sabatini S. 2010. Analysis of Root Meristem Size Development. Plant Developmental Biology: Methods and Protocols 655: 177–187.

Persson DP, Chen A, Aarts MGM, Salt DE, Schjoerring JK, Husted S. 2016. Multi-element bioimaging of Arabidopsis thaliana roots. Plant Physiology 172: 835–847.

Petersen K, Fiil BK, Mundy J, Petersen M. 2008. Downstream targets of WRKY33. Plant Signaling and Behavior 3: 1033–1034.

Rajniak J, Giehl RFH, Chang E, Murgia I, Von Wirén N, Sattely ES. 2018. Biosynthesis of redox-active metabolites in response to iron deficiency in plants. Nature Chemical Biology 14: 442–450.

Ravet K, Touraine B, Kim SA, Cellier F, Thomine S, Guerinot M Lou, Briat JF, Gaymard F. 2009. Post-translational regulation of AtFER2 ferritin in response to intracellular iron trafficking during fruit development in Arabidopsis. Molecular Plant 2: 1095–1106.

Remans T, Opdenakker K, Guisez Y, Carleer R, Schat H, Vangronsveld J, Cuypers A. 2012. Exposure of Arabidopsis thaliana to excess Zn reveals a Zn-specific oxidative stress signature. Environmental and Experimental Botany 84: 61–71.

Reyt G, Boudouf S, Boucherez J, Gaymard F, Briat JF. 2015. Iron- and ferritin-dependent reactive oxygen species distribution: Impact on Arabidopsis root system architecture. Molecular Plant 8: 439– 453.

Riaz N, Guerinot M Lou. 2021. All together now: Regulation of the iron deficiency response. Journal of Experimental Botany 72: 2045–2055.

Ricachenevsky FK, Menguer PK, Sperotto RA, Fett JP. 2015. Got to hide your Zn away: Molecular control of Zn accumulation and biotechnological applications. Plant Science 236: 1–17.

Richtmann L, Thiébaut N, Sarthou M, Ranjan A, Boutet S, Hanikenne M, Verbruggen N, Clemens S. A multi-omics analysis of Arabidopsis thaliana root tips under Cd exposure: Insights into HY5’s role in limiting accumulation. Submitted as companion paper to New Phytologist.

Schaller GE, Bishopp A, Kieber JJ. 2015. The yin-yang of hormones: Cytokinin and auxin interactions in plant development. Plant Cell 27: 44–63.

Scheepers M, Spielmann J, Boulanger M, Carnol M, Bosman B, De Pauw E, Goormaghtigh E, Motte P, Hanikenne M. 2020. Intertwined metal homeostasis, oxidative and biotic stress responses in the Arabidopsis frd3 mutant. Plant Journal 102: 34–52.

Schwarz B, Bauer P. 2020. FIT, a regulatory hub for iron deficiency and stress signaling in roots, and FIT-dependent and -independent gene signatures. Journal of Experimental Botany 71: 1694–1705.

Shahan R, Hsu CW, Nolan TM, Cole BJ, Taylor IW, Greenstreet L, Zhang S, Afanassiev A, Vlot AHC, Schiebinger G, et al. 2022. A single-cell Arabidopsis root atlas reveals developmental trajectories in wild-type and cell identity mutants. Developmental Cell 57: 543–560.e9.

Shanmugam V, Lo JC, Wu CL, Wang SL, Lai CC, Connolly EL, Huang JL, Yeh KC. 2011. Differential expression and regulation of iron-regulated metal transporters in Arabidopsis halleri and Arabidopsis thaliana - the role in zinc tolerance. New Phytologist 190: 125–137.

Sinclair SA, Krämer U. 2012. The zinc homeostasis network of land plants. BBA - Molecular Cell Research 1823: 1553–1567.

Sinclair SA, Senger T, Talke IN, Cobbett CS, Haydon MJ, Krämer U. 2018. Systemic upregulation of mtp2-and hma2-mediated zn partitioning to the shoot supplements local zn deficiency responses[open]. Plant Cell 30: 2463–2479.

Sinclair SA, Sherson SM, Jarvis R, Camakaris J, Cobbett CS. 2007. The use of the zinc-fluorophore, Zinpyr-1, in the study of zinc homeostasis in Arabidopsis roots. New Phytol 174: 39–45.

De Smet S, Cuypers A, Vangronsveld J, Remans T. 2015. Gene Networks Involved in Hormonal Control of Root Development in Arabidopsis thaliana: A Framework for Studying Its Disturbance by Metal Stress. International Journal of Molecular Sciences 16: 19195–19224.

Sofo A, Bochicchio R, Amato M, Rendina N, Vitti A, Nuzzaci M, Altamura MM, Falasca G, Rovere F Della, Scopa A. 2017. Plant architecture, auxin homeostasis and phenol content in Arabidopsis thaliana grown in cadmium- and zinc-enriched media. Journal of Plant Physiology 216: 174–180.

Somssich M, Khan GA, Persson S. 2016. Cell Wall Heterogeneity in Root Development of Arabidopsis. Frontiers in Plant Science 07: 1–11.

Spielmann J, Cointry V, Devime F, Ravanel S, Neveu J, Vert G. 2022. Differential metal sensing and metal-dependent degradation of the broad spectrum root metal transporter IRT1. Plant Journal: 1252– 1265.

Spielmann J, Schloesser M, Hanikenne M. 2024. Reduced expression of bZIP19 and bZIP23 increases zinc and cadmium accumulation in Arabidopsis halleri. Plant, Cell & Environment 47: 2093–2108.

Spielmann J, Vert G. 2021. The many facets of protein ubiquitination and degradation in plant root iron-deficiency responses. Journal of Experimental Botany 72: 2071–2082.

Stanton C, Rodríguez-Celma J, Krämer U, Sanders D, Balk J. 2023. BRUTUS-LIKE (BTSL) E3 ligase-mediated fine-tuning of Fe regulation negatively affects Zn tolerance of Arabidopsis. Journal of Experimental Botany 74: 5767–5782.

Stanton C, Sanders D, Krämer U, Podar D. 2022. Zinc in plants: Integrating homeostasis and biofortification. Molecular Plant 15: 65–85.

Stortenbeker N, Bemer M. 2019. The SAUR gene family: the plant’s toolbox for adaptation of growth and development. Journal of Experimental Botany 70: 17–27.

Stotz HU, Sawada Y, Shimada Y, Hirai MY, Sasaki E, Krischke M, Brown PD, Saito K, Kamiya Y. 2011. Role of camalexin, indole glucosinolates, and side chain modification of glucosinolate-derived isothiocyanates in defense of Arabidopsis against Sclerotinia sclerotiorum. Plant Journal 67: 81–93.

Talke IN, Hanikenne M, Krämer U. 2006. Zinc-dependent global transcriptional control, transcriptional deregulation, and higher gene copy number for genes in metal homeostasis of the hyperaccumulator Arabidopsis halleri. Plant Physiology 142: 148–167.

Tennstedt P, Peisker D, Böttcher C, Trampczynska A, Clemens S. 2009. Phytochelatin synthesis is essential for the detoxification of excess zinc and contributes significantly to the accumulation of zinc. Plant Physiology 149: 938–948.

Tewes LJ, Stolpe C, Kerim A, Krämer U, Müller C. 2018. Metal hyperaccumulation in the Brassicaceae species Arabidopsis halleri reduces camalexin induction after fungal pathogen attack. Environmental and Experimental Botany 153: 120–126.

Thiébaut N, Hanikenne M. 2022. Zinc deficiency responses: bridging the gap between Arabidopsis and dicotyledonous crops. Journal of Experimental Botany 73: 1699–1716.

Thomma BPHJ, Nelissen I, Eggermont K, Broekaert WF. 1999. Deficiency in phytoalexin production causes enhanced susceptibility of Arabidopsis thaliana to the fungus Alternaria brassicicola. Plant Journal 19: 163–171.

Tsai HH, Rodríguez-Celma J, Lan P, Wu YC, Vélez-Bermúdez IC, Schmidt W. 2018. Scopoletin 8-hydroxylase-mediated fraxetin production is crucial for iron mobilization. Plant Physiology 177: 194– 207.

Tsuji J, Jackson EP, Gage DA, Hammerschmidt R, Somerville SC. 1992. Phytoalexin accumulation in Arabidopsis thaliana during the hypersensitive reaction to Pseudomonas syringae pv syringae. Plant Physiology 98: 1304–1309.

Tsukagoshi H, Busch W, Benfey PN. 2010. Transcriptional regulation of ROS controls transition from proliferation to differentiation in the root. Cell 143: 606–616.

Vélez-Bermúdez IC, Schmidt W. 2023. Iron sensing in plants. Frontiers in Plant Science 14: 1–7.

Vert G, Barberon M, Zelazny E, Séguéla M, Briat JF, Curie C. 2009. Arabidopsis IRT2 cooperates with the high-affinity iron uptake system to maintain iron homeostasis in root epidermal cells. Planta 229: 1171–1179.

Vestenaa MW, Husted S, Minutello F, Persson DP. 2024. Endodermal suberin restricts root leakage of cesium: a suitable tracer for potassium. Physiologia Plantarum 176: 1–14.

Vik D, Mitarai N, Wulff N, Halkier BA, Burow M. 2018. Dynamic modeling of indole glucosinolate hydrolysis and its impact on auxin signaling. Frontiers in Plant Science 9: 1–16.

Wang J, Moeen-ud-din M, Yang S. 2021. Dose-dependent responses of Arabidopsis thaliana to zinc are mediated by auxin homeostasis and transport. Environmental and Experimental Botany 189: 104554.

Won SK, Lee YJ, Lee HY, Heo YK, Cho M, Cho HT. 2009. cis-element- and transcriptome-based screening of root hair-specific genes and their functional characterization in Arabidopsis. Plant Physiology 150: 1459–1473.

Yang W, Wightman R, Meyerowitz EM. 2017. Cell Cycle Control by Nuclear Sequestration of CDC20 and CDH1 mRNA in Plant Stem Cells. Molecular Cell 68: 1108–1119.e3.

Yuan HM, Huang X. 2016. Inhibition of root meristem growth by cadmium involves nitric oxide-mediated repression of auxin accumulation and signalling in Arabidopsis. Plant Cell and Environment 39: 120–135.

Zamioudis C, Korteland J, Van Pelt JA, Van Hamersveld M, Dombrowski N, Bai Y, Hanson J, Van Verk MC, Ling HQ, Schulze-Lefert P, et al. 2015. Rhizobacterial volatiles and photosynthesis-related signals coordinate MYB72 expression in Arabidopsis roots during onset of induced systemic resistance and iron-deficiency responses. The Plant Journal 84: 309–322.

Zhang P, Sun L, Qin J, Wan J, Wang R, Li S, Xu J. 2018. cGMP is involved in Zn tolerance through the modulation of auxin redistribution in root tips. Environmental and Experimental Botany 147: 22–30.

Zhong K, Zhang P, Wei X, Platre MP, He W, Zhang L, Małolepszy A, Cao M, Hu S, Tang S, et al. 2024. Natural variation of TBR confers plant zinc toxicity tolerance through root cell wall pectin methylesterification. Nature Communications 15: 1–14.

Zluhan-Martínez E, López-Ruíz BA, García-Gómez ML, García-Ponce B, de la Paz Sánchez M, Álvarez-Buylla ER, Garay-Arroyo A. 2021. Integrative Roles of Phytohormones on Cell Proliferation, Elongation and Differentiation in the Arabidopsis thaliana Primary Root. Frontiers in Plant Science 12: 1–20.

