## Supplementary material for "Specific redox and iron homeostasis responses in the root tip of Arabidopsis upon zinc excess": Thiebaut_Supporting information

**Table of Contents**

Methods S1 2-9

Figures S1-S17 10-29

Tables S1-S7 (Captions) 30

References 31-33

**Methods S1**

This section presents detailed Materials and Methods.

**Plant material and growth conditions**

*A. thaliana* (Col-0) was used in all experiments. The *pad3-1* (*phytoalexin deficient*, (Glazebrook *et al.*, 1997), *pad3-3* (SALK_026585C), *pad3-4* (SALKseq_059784.1) were obtained from NASC (Nottingham Arabidopsis Stock Centre) and used as homozygote lines.

Plants were grown vertically on agar plates with Hoagland medium [1.5 mM $Ca(NO_3)_2$, 0.28 mM $KH_2PO_4$, 0.75 mM $MgSO_4$, 1.25 mM $KNO_3$, 0.5 µM $CuSO_4$, 1 µM $ZnSO_4$, 5 µM $MnSO_4$, 25 µM $H_3BO_3$, 0.1 µM $Na_2MoO_4$, 50 µM KCl, 10 µM Fe-HBED, 0.5 g/L MES, 1% (w/v) sucrose, 0.8 % (w/v) agar (M-type, Sigma), pH 5.7] in BrightBoy GroBanks (CLF Plant Climatics) under long day conditions (16 h light, 21°C, 100 $\mu mol.m^{-2}.s^{-1}$ / 8 h dark, 18°C). For the Laser Ablation ICP-MS experiment, seedlings were grown under slightly different conditions [16 h light, 22°C, 125 $\mu mol.m^{-2}.s^{-1}$ / 8 h dark, 20°C, in a Percival chamber (CLF Plant Climatics)].

Seeds were surface sterilized by rinsing 3 min with 70 % ethanol and 2 min with 2 % bleach. After 3 washing steps with sterile water, the seeds were suspended in 0,1 % (w/v) sterile agar. For most experiments, seeds were sown on control agar plates on nylon meshes (200 µm aperture, Prosep filters) and stratified (4°C, 2 days) (Fig. **S1**). After 7 days, seedlings were transferred to either new control plates (1 µM Zn) or Zn excess plates (200 µM Zn), and treated for 24 hours, 48 hours or 7 days (Fig. **S1**). The experimental design combining germination on control medium then transfer to a high Zn concentration (200 µM) was selected based on preliminary experiments, meeting several requirements: it enabled (i) initial seedling establishment in a non-toxic condition, reduced growth shortly after transfer, (ii) the capture of the initial changes in the root tip (RT) upon Zn exposure, (iii) survival of the seedlings. For the *pad3* mutant characterization, seeds were sown directly on control (1 µM Zn) or treated (75 µM Zn) Hoagland media, supplemented or not with 250 nM of camalexin (Sigma Aldrich), and grown for 11 days. This simplified the handling of the experiments. Using lower Zn concentrations for a longer time achieved an overall similar growth reduction than in transfer experiments.

Growth assays under Fe-Zn excess conditions (Fig. **S16**) were carried out in the same conditions as for the other *pad3* mutant experiments, except that seedlings were grown on control (1 µM Zn, 10 µM Fe), or upon Fe (1 µM Zn, 70 µM Fe), Zn (75 µM Zn, 10 µM Fe) or Fe-Zn (75 µM Zn, 70 µM Fe) excesses.

Plants used for the RT-qPCR to investigate the effect of Fe deficiency on *PAD3* expression were hydroponically grown in control liquid Hoagland medium for 3 weeks (short days, 8h light, 22°C, 100 $\mu mol.m^{-2}.s^{-1}$ / 16h dark, 20°C). They were then transferred for 3 weeks into control (10 µM Fe) or treatment media (0 µM Fe).

A description of the experimental replication levels is presented in each figure legend.

**Root growth and RAM phenotyping**

At three independent time-points (after the transfer to fresh plates at day 7, and after 24h and 48h of treatment), the root apex was marked at the back of the plate for root growth measurement (Fig. 1A). Plates were scanned with an EPSON V3300 scanner after 7 days of treatment and roots were measured with FIJI (https://imagej.net/Fiji). To measure the root apical meristem (RAM) size, seedlings treated for 24h and 48h were rinsed 30 seconds in distilled water, then stained in the dark for 2 minutes in propidium iodide (33.4 mg/L) and rinsed again in distilled water. Roots were mounted on glass slides and visualized with a SP2 TCS AOBS confocal laser-scanning inverted microscope (Leica, Mannheim, Germany) with excitation at 543 nm and emission between 600 and 730 nm. The pictures of the propidium iodide-stained roots were then used to quantify several morphological features using FIJI, as follows. The length of the meristematic zone was assessed as the distance between the first elongated cortex cell and the quiescent center (QC) (Perilli & Sabatini, 2010). The number of cortex cells in this zone was simultaneously assessed and a mean cortex cell size in RAM was calculated by dividing the length (in µm) by the number of cortex cells for each root. To estimate the size of the elongation zone (EZ), the distance between the root apex and the first root hair was measured for each root.

**Ionome profiling**

For each sample, seedlings from 48 plates, containing each 16 to 18 seedlings, were harvested after 24h or 48h of treatment, pooled and desorbed prior to ionome analysis as follows. The seedlings were washed twice in a desorption solution (5 mM $CaCl_2$, in 1 mM MES, pH5.7) for 10 min at 4°C and twice in distilled water (5 min, 4°C). About 2-3 mm long root tips (RT) were then separated from the remaining root system (RR) with a clean scalpel blade on a clean surface. Dry root material was weighed into polytetrafluoroethylene digestion tubes and concentrated nitric acid [0.5 ml, 67-69 % (v/v)] was added to each tube. After 4 h incubation, samples were digested under pressure using a high-performance microwave reactor (Ultraclave 4, MLS). Digested samples were transferred to Greiner centrifuge tubes and diluted with de-ionized water. Elemental analysis was carried out using ICP-MS (Induced Coupled Plasma-Mass spectrometry analysis) [Sector Field High Resolution (HR)-ICP-MS, ELEMENT 2, Thermo Fisher Scientific]. For sample introduction a SC-2 DX Autosampler (ESI, Elemental Scientific) was used. A 6 points external calibration curve was set from a certified multiple standard solution (Bernd Kraft, Duisburg, Germany). Elements rhodium (Rh) and germanium (Ge) were infused online and used as internal standards.

For the *pad3* mutant characterization, samples were harvested after 11 days of growth under control, zinc excess (75 µM), and/or camalexin supplementation (250 nM) conditions on Hoagland agar plates.

Shoots and roots from 60 to 90 seedlings per sample were then pooled. The pooled roots were rinsed twice in a desorption solution (5 mM $CaCl_2$, in 1 mM MES, pH 5.7) for 10 min at 4°C and twice in ultrapure Milli-Q water (Millipore) (5 min, 4°C). The pooled shoots were rinsed 5 min in ultrapure Milli-Q water (Millipore). Dried root and shoot samples (48h at 55°C) were weighed and digested in 1.5 mL of 65% $HNO_3$ by gradually heating (15 minutes at 45°C, 15 minutes at 65°C) up to 95°C for a 60-minute incubation. The digested samples were diluted to a final volume of 5 mL before analysis by ICP-OES (Inductively Coupled Plasma Optical Emission Spectroscopy) using a 5110 SVDV ICP-OES device (Agilent).

**Laser Ablation ICP-MS**

Roots were harvested and two regions of interest (ROI) along the primary roots were prepared: RAM (~200 µm from the tip of the columella) and differentiated roots (at a distance from the apex corresponding to 50 and 60% of root length). ROI of primary roots from several seedlings were aligned in a structure called "root bouquet" (Fig. **S2**). The portion of the bouquet containing either the aligned root tips or ROI of differentiated roots was dipped into liquid paraffin (42°C), dried for 3 s, re-dipped in liquid paraffin and placed in optimal cutting temperature media (OCT) in a rubber mold, before freezing in liquid nitrogen. At least seven 14 µm-thick perpendicular sections of the samples were cut in a cryotome (Leica CM050S, St. Gallen, Switzerland; precooled to −25°C), and transferred to microscopy glass slides, using Cryofilm 2C (SectionLab) as described in (Persson *et al.*, 2016). The first and last sections were used for fresh microscopy pictures, and the five middle sections were dried at -15°C (Fig. S2). The best dried section was photographed and selected for laser ablation ICP-MS (Fig. S2). Two bouquets were analyzed, as technical duplicates, for each condition and root part.

The Laser Ablation ICP MS system was composed of an Iridia Laser Ablation system (Teledyne CETAC Technologies, United Kingdom), coupled to a ICP-QQQ-MS 8900 instrument (Agilent, United Kingdom). Samples were digested at a spot size of 4 µm diameter, at 200 Hz, with a fluence of 0.5 $J.cm^{-2}$. For the detection, three elements were analyzed; $^{39}K$, $^{56}Fe$ and $^{66}Zn$ with integration times of 0.001, 0.01 and 0.07 s, respectively. The HDIP software was used for data analysis (Teledyne CETAC Technologies). Signal intensity across the root was taken by selecting a 20 µm-wide zone across all cell layers, as represented in Supplemental Fig. **S2d-e**, and used to represent signal over distance. With help of the HDIP software (Teledyne Photon Machine), the distance was cropped manually to each side of the roots, to exactly the external side of the epidermis. This distance was normalized to radius=1, and centered around 0.

**RNA-Seq analysis**

For RNA sequencing, a total of 48 plates containing 16-18 seedlings each were harvested per independent replicate per sample. RT samples were obtained by cutting at ~2-3 mm from the apex conducted, rapidly collected and frozen in liquid nitrogen (Fig. **S1**). RR tissues were also collected and frozen. Total RNA isolation was conducted with a Maxwell 16 device (Promega) using the Maxwell RSC Plant RNA Kit (Promega). RNA quality and concentration were measured with an Agilent 2100 Bioanalyzer Expert using the Agilent Nano 6000 Kit. Library preparation was performed with the TruSeq Stranded mRNA Sample Preparation Kit (Illumina), using 1 µg of total RNA as starting material. Library quality was assessed with a QIAxcel screening kit (Qiagen). Quantification, dilution and pooling of the libraries were done using the KAPA SYBR® FAST Universal qPCR Kit (Roche) and an Applied Biosystems 7900HT Real-Time PCR system. Sequencing (PE 2x150 nt) was done at the GIGA Genomics platform (University of Liège, Belgium) on a NovaSeq6000 device (Illumina) with standard workflow to obtain ~10 Mio reads/sample.

Data quality assessment and trimming were done with FastQC & Trimmomatic (Bolger *et al.*, 2014) with the following parameters: leading = 30, trailing =30, slidingwindow = 10:30, crop = 98, minlen =98. HiSAT2 was used for mapping on the *A. thaliana* genome [TAIR10 genome assembly and Araport11 annotation (Cheng *et al.*, 2017)]. Read counting was done with htseq-count (Anders *et al.*, 2015) and differential expression analysis with DESeq2 package (Love *et al.*, 2014). Calculation of differential expression between Zn excess samples and the corresponding control samples was realized with the following function and parameters: DESeq2::results(Def_DESeq, contrast, lfcThreshold = 1, alpha = 0.05), followed by padj < 0.05 and either $\log_2$FoldChange > 1 or $\log_2$FoldChange < -1 for selection of upregulated and downregulated genes, respectively.

GO enrichment analysis for biological processes was conducted with the Panther Database (https://pantherdb.org/) in September 2022, against *A. thaliana* annotated genes, with Bonferroni-correction with pval < 0.05. For graphical representation, GO represented by only one DEG were omitted, as well as two additional GO annotations which were too wide to provide useful information: *cellular process* (GO:0009987) and *biological_process* (GO:0008150).

**RT qPCR**

Harvesting was carried out by separating the roots and shoots of 11-day-old seedlings, freezing them in liquid nitrogen and cold-grinding them twice using a ball mill at 30 Hz for 60 s (Retsch, Germany). Samples were collected in triplicates of 40 to 50 seedlings. RNA was extracted using the NucleoSpin® RNA Plant Kit (Machery-Nagel, Germany), following the kit instructions. RNA quality and quantity were assessed by spectrophotometric analysis, and 1 µg was used for cDNA synthesis, using the RevertAid First Strand cDNA Synthesis Kit (Thermo Fisher Scientific, USA), according to the manufacturer instructions. RT-qPCR were performed on a QuantStudio™ 5 (Thermo Fisher Scientific, USA). Each reaction contained 5 µL of Takyon™ No ROX SYBR 2X MasterMix (Eurogentec, Belgium), 1 µL of 2.5 µM forward-reverse primer mix (Table **S1**) and 4 µL of 1:50 diluted cDNA. For amplification, cDNA was preincubated 2 min at 50°C, then 2 min at 95°C, followed by 40 cycles of 15 s at 95°C and 1 min at 60°C, and a final step of 15 s at 95°C, 1 min at 60 °C and 15 s at 95°C to determine dissociation curves. Data were analyzed using the Design & Analysis Software (v. 2.8.0, Thermo Fisher Scientific, USA), and qbase+ (v. 3.4.1, Biogazelle, Belgium). The analysis was done with three technical replicates for each sample/primer combination, and *UBIQUITIN10* (UBQ10), *EF1alpha* and *AT1G58050* were used as reference genes for data normalization.

**Untargeted metabolomic analysis**

For untargeted metabolomic analysis (Boutet *et al.*, 2022), plants were grown as described above. 48h after transferring seedlings to control or Zn excess plates, RT and RR samples were collected by cutting and collecting similar sized pieces of ~2-3 mm of RT and RR (~mid-distance from the apex).

*Extraction of polar and semipolar* primary and specialized metabolites***.*** Metabolites were extracted from Arabidopsis root tissues by 2*1.8 mL of MeOH:$H_2O$:Acetone:TFA (30/28/42/0.1 V/V/V/V) and 500 ng of apigenin (Api, used as internal standard) were added to each sample. the supernatants were collected and dried using a speed vac before being resuspended in 200 µL of water and 10% acetonitrile for LC-MS/MS analysis.

*Specialized metabolite extraction and UPLC-MS/MS analysis*. Untargeted metabolomics data were acquired using a UHPLC system (Ultimate 3000 Thermo) coupled to quadrupole time of flight mass spectrometer (Q-Tof Impact II Bruker Daltonics, Bremen, Germany). For untargeted polar and semipolar metabolomic analysis, a Nucleoshell RP 18 plus reversed-phase column (2 x 100 mm, 2.7 µm; Macherey-Nagel) was used for chromatographic separation. The mobile phases used for the chromatographic separation were (A) 0.1% formic acid in $H_2O$ and (B) 0.1% formic acid in acetonitrile. The flow rate was of 400 µL/min and the following gradient was used: 95% of A for 1-min, followed by a linear gradient from 95% A to 80% A from 1 to 3-min, then a linear gradient from 80% A to 75% A from 3 to 8-min, a linear gradient from 75% A to 40% A from 8 to 20-min. 0% of A was hold until 24-min, followed by a linear gradient from 0% A to 95% A from 24 to 27-min. Finally, the column was washed by 30% A at for 3.5-min then re-equilibrated for 3.5-min (35-min total run time). Data-dependent acquisition (DDA) methods were used for mass spectrometer data in positive and negative ESI modes using the following parameters: capillary voltage, 4.5kV; nebulizer gas flow, 2.1 bar; dry gas flow, 6 L/min; drying gas in the heated electrospray source temperature, 140°C. Samples were analyzed at 8 Hz with a mass range of 100 to 1500 m/z. Stepping acquisition parameters were created to improve the fragmentation profile with a collision RF from 200 to 700 Vpp, a transfer time from 20 to 70 µs and collision energy from 20 to 40 eV. Each cycle included a MS fullscan and 5 MS/MS CID on the 5 main ions of the previous MS spectrum.

*Metabolomic data processing.* The .d data files (Bruker Daltonics, Bremen, Germany) were converted to .mzXML format using DataAnalysis from Bruker Daltonics. mzXML data processing, mass detection, chromatogram building, deconvolution, samples alignment, and data export, were performed using the MZmine 2.52 software (http://mzmine.github.io/) for both positive and negative data files. The ADAP chromatogram builder (Myers *et al.*, 2017) method was used with a minimum group size of scan 5, a group intensity threshold of 5000, a minimum highest intensity of 5000 and m/z tolerance of 10 ppm. Deconvolution was performed with the ADAP wavelets algorithm using the following setting: S/N threshold 8, peak duration range = 0.01-2.1 min retention time wavelet range 0.0-0.05 min, MS2 scan were paired using a m/z tolerance range of 0.2 Da and retention time tolerance of 0.2 min. Then, isotopic peak grouper algorithm was used with a m/z tolerance of 10 ppm and retention time tolerance of 0.5. All the peaks were filtered using feature list row filter keeping only peaks with MS2 scan. The alignment of samples was performed using the join aligner with an m/z tolerance of 10 ppm, a weight for m/z and retention time at 1, a retention time tolerance of 0.5 min. First research in library with Mzmine was done with identification module and “custom database search” to begin the annotation with our library, currently containing 103 annotations (retention time and m/z) in positive mode and 67 in negative mode, with retention time tolerance of 0.3 min and m/z tolerance of 0.0025 Da or 6 ppm.

*Molecular networking.* Molecular networks were generated with MetGem software (Olivon *et al.*, 2018; https://metgem.github.io) using the .mgf and .csv files obtained with MZmine2 analysis. The molecular network was optimized for the ESI+ and ESI- datasets and different cosine similarity score thresholds were tested. ESI- and ESI+ molecular networks were generated using cosine score thresholds of 0.8, in positive mode and negative mode. Optimal parameters that allowed a good cluster separation among metabolites were set up.

*Metabolite annotations.* Metabolite annotations were performed in four consecutive steps. First, the obtained RT and m/z data of each feature were compared with our home-made library containing more than 150 standards or experimental common features (retention time, m/z). Second, the ESI- and ESI+ metabolomic data used for molecular network analyses were searched against the available MS2 spectral libraries (Massbank NA, GNPS Public Spectral Library, NIST14 Tandem, NIH Natural Product and MS-Dial), with absolute m/z tolerance of 0.02, 4 minimum matched peaks and minimal cosine score of 0.65. Third, not-annotated metabolites that belong to molecular network clusters containing annotated metabolites from step 1 and 2 were assigned to the same chemical family. Finally, for metabolites that had no or unclear annotation, the Sirius software (https://bio.informatik.uni-jena.de/software/sirius/) was used. Sirius is based on machine learning techniques that use available chemical structures with the CANOPUS module (Dührkop *et al.*, 2021) and MS/MS data from chemical databanks to propose structures of unknown compounds.

*Metabolite normalization.* Considering (i) the small size of the samples and the difficulty of weighting them, and (ii) the very different total abundances of metabolites between the two root parts (6 times more abundant in RR than RT, Fig. **S17b**), two methods of data normalization were considered : (i) normalization according to the number of collected root parts (Fig. **S17a,b**), and (ii) normalization according to the total metabolite abundance in each sample (Fig. **6a**). The two methods differently impacted how the variance was partitioned among root part (78% vs 57%, respectively) and treatment (13% vs 18%, respectively) (Fig. **S17a,** Fig. **6a**). As this method provides consistent total metabolite abundance across samples, which facilitates reliable comparisons, and is commonly used in the literature (Holmes *et al.*, 2008; Di Guida *et al.*, 2016; Pang *et al.*, 2022), data were normalized by the total count of control samples, enabling a detailed analysis of the impact of the treatment in each root part.

**Targeted camalexin analysis**

Camalexin extraction was adapted from Tewes *et al.* (2018). Upon harvest, roots and shoots from 11 day-old Col-0 and *pad3-3* seedlings were separated, then weighted (100 to 500 mg) and frozen in liquid nitrogen. Plant material was then frozen ground using a ball mill at 30 Hz for 60 s, twice (Retsch, Germany). A first camalexin extraction was performed using 400 to 500 µL of 80 % (v/v) HPLC grade methanol (Sigma Aldrich, USA). Samples were homogenized and incubated for 10 min at 60°C and 1000 rpm in a reaction block (Thermomixer, Eppendorf, Germany). Samples were centrifuged at room temperature for 5 min at 17 000 g. The supernatant was collected in a new tube and the extraction operation was repeated a second time. The two joined supernatants were dried under reduced pressure with a RVC 2-33 CDplus (Christ, Germany). Samples were resuspended in 200 to 1000 µL of 80 % methanol, filtered through a polytetrafluoroethylene (PTFE) membrane filter and analyzed using Ultra-Performance Liquid Chromatography–High-Resolution Mass Spectrometry (UPLC-HRMS).

The UPLC-HRMS analyses were performed on a Waters ACQUITY Ultra Performance LC system equipped with a Waters Synapt XS Q-TOF. A 5.0 µl sample solution was injected for each run. Chromatographic separation was performed with a flow of 0.5 ml/min, at 40°C, using the 1.7 µm ACQUITY UPLC BEH C18 Column (2.1 mm X 50 mm dimension, 130 Å pores). The mobile phase consisted of water (containing 0.1% formic acid – solvent A) and acetonitrile (containing 0.1% formic acid – solvent B). The elution gradient was 0 - 1 min, 5% B; 1.0 - 25.0 min, 5 - 100% B; 25.0 - 28.0 min, 100% B; 28.0 - 28.5 min, 100.0 - 5.0% B; 28.5- 30.0 min, 5.0% B. UV spectra was recorded with the built-in ACQUITY UPLC Photodiode Array (PDA) detector with a wavelength ranging from 190 to 500 nm, resolution of 1.2 nm and a sampling rate of 20 points/sec. Mass spectrometry analyses were performed on positive electrospray polarity with a source temperature of 150°C. The capillary and cone voltage were set at 0.5 kV and 70 V, respectively. An MSe continuum function in resolution mode with an extended dynamic range, a scan range of 50 to 1200 m/z and a scan time of 0.15 seconds was used. For the high energy (MS2) data collection a collision energy ramp from 20 to 70 V was set for the transfer cell. All mass spectrometry analyses were acquired using Leucine-enkephalin solution (100 ng/ml in $H_2O$/ACN 50:50) as the internal calibrant. All LC-MS data analysis was performed on Masslynx V 4.2 SCN1015. Lockspray correction was performed based on the calculated monoisotopic mass of the M+H+ ion of Leucine-enkephalin (556.2766).

Camalexin was identified based on the retention time of an external standard of commercial camalexin (Sigma-Aldrich), and the identity was confirmed based on the m/z value of camalexin's [M+H] adduct detected on the mass spectra (mass error lower than 5 ppm). For camalexin quantification, Extracted Ion Chromatograms (EIC) for camalexin's [M+H] ion was generated for seven standard camalexin solutions (0.0 µM, 0.05 µM, 0.2 µM, 0.5 µM, 1.0 µM, 2.0 µM, 3.0 µM, 5.0 µM) from which peak area was calculated and used to construct a calibration curve. Concentration was then reported per gram of fresh weight.

**Supporting Information Figures**

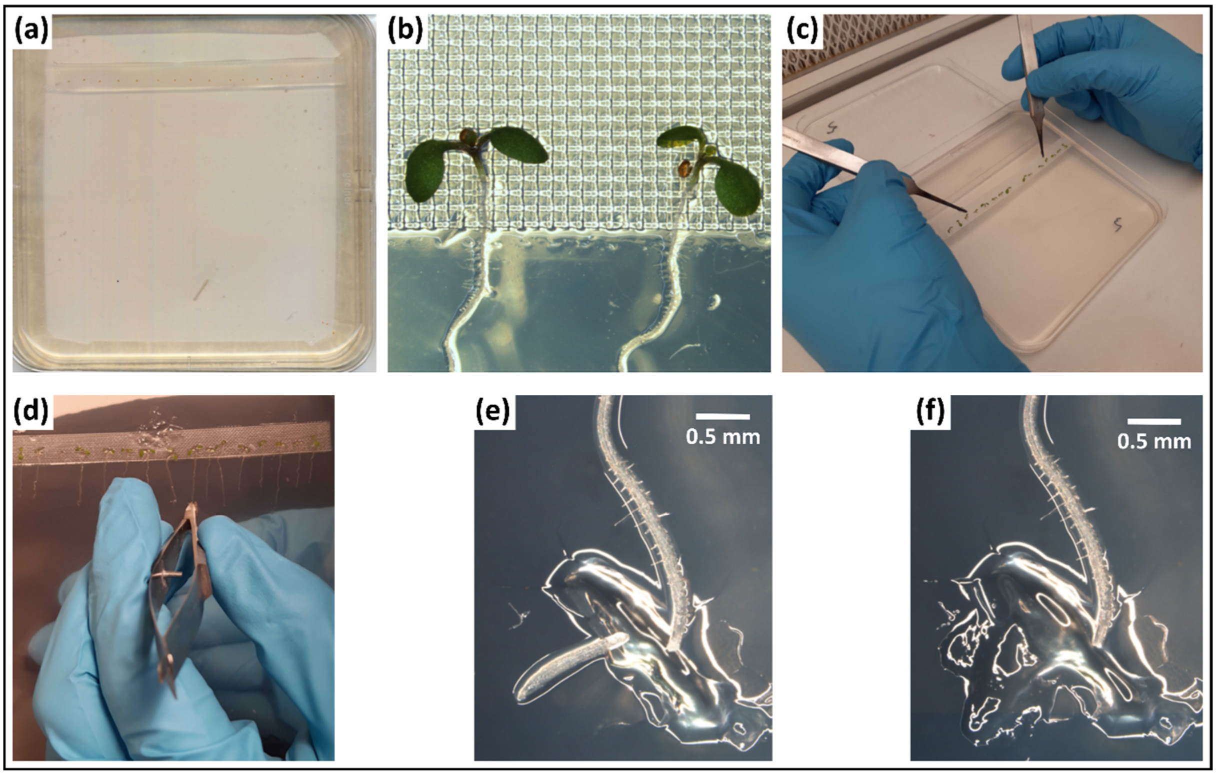


**Figure S1. Illustration of growth conditions and sampling of root tip and remaining roots for the RNA-Seq and ICP-MS analyses. a.** Seeds sown on nylon mesh in a vertical plate. **b.** Germinated seedlings on the nylon mesh. **c.** Nylon mesh and seedling transfer to fresh media. **d.** On agar plate microdissection of the Root Tip (RT). **e.** Freshly cut RT on the agar plate. **f.** Remaining root after RT collection on agar media.

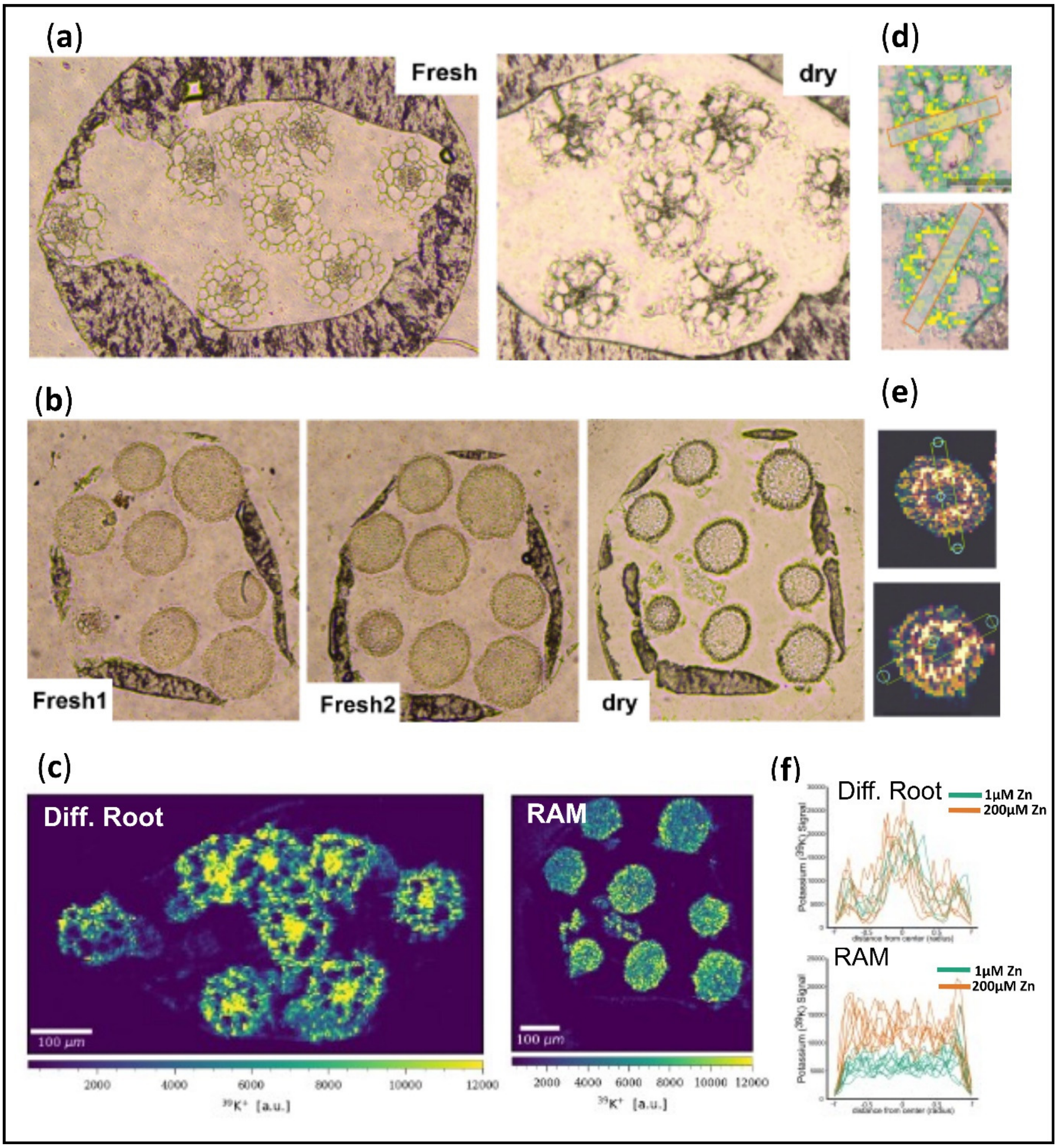


**Figure S2. Processing of Laser Ablation ICP-MS samples. a-b.** Microscopic pictures of bouquet sections of differentiated roots (**a**) and RAM (**b**) samples embedded in paraffin. **c.** Representative potassium $^{39}K$ signal, used as a control, in differentiated root and RAM bouquets. **d-e.** Illustration of signal intensity collection within a region on interest, marked by a rectangle, across differentiated roots (**d**) and RAM (**e**). **f.** $^{39}K$ signal in differentiated roots (RR) and RAM (RT) bouquets. Each graph contains data for 8 differentiated root or RAM sections per condition, coming from two distinct root bouquets (see Methods S1 for details).

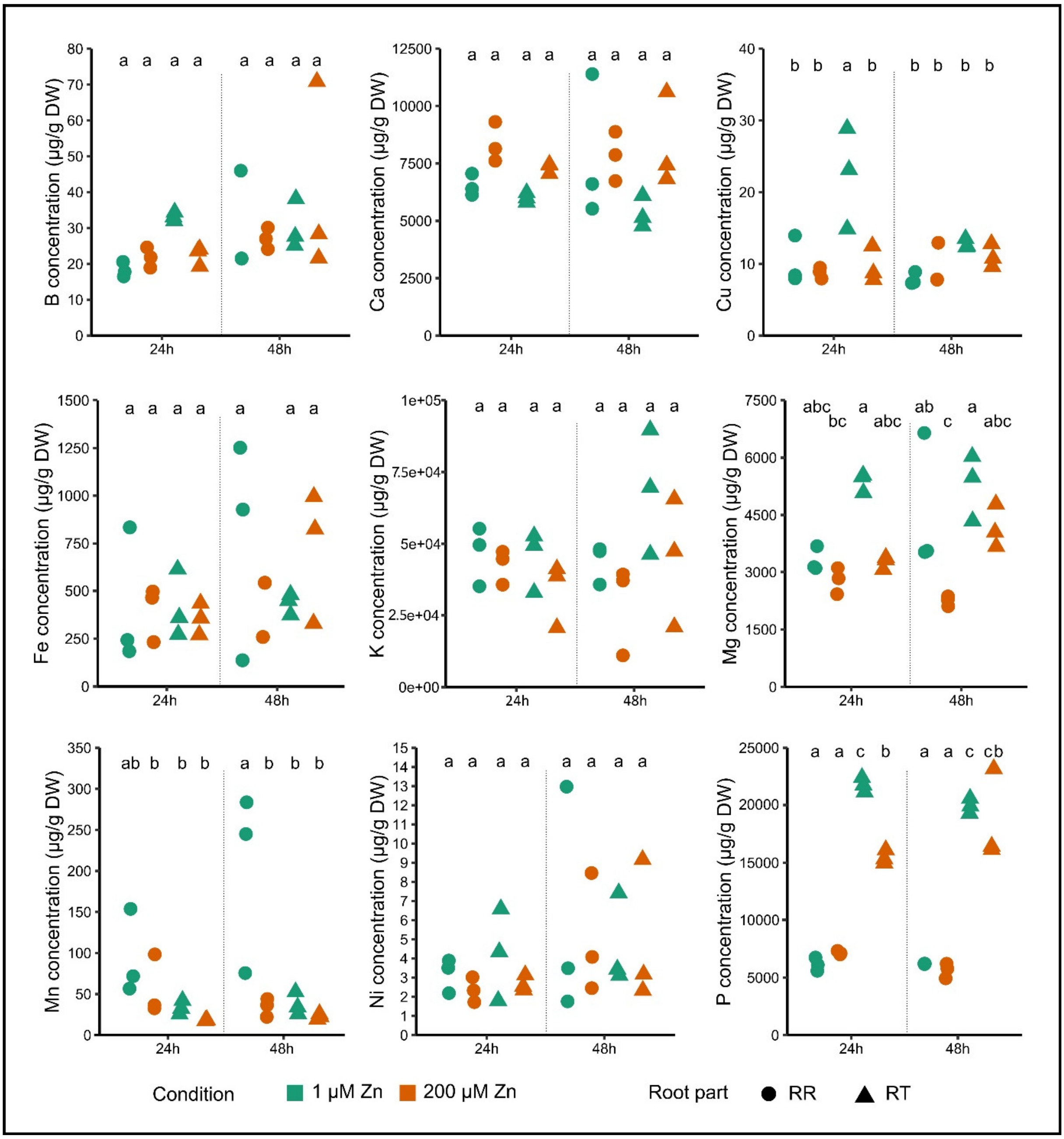


**Figure S3. Ionome profiling of the root tip and the remaining root system upon Zn excess in Arabidopsis.** Seedlings were germinated and grown for one week in control agar plates (1 µM Zn), then transferred for 24h or 48h onto new control or Zn excess (200 µM Zn) plates. Root tip (RT) and remaining root (RR) samples were then analyzed by ICP-MS. B, Ca, Cu, Fe, K, Mg, Mn, Ni and P concentrations are presented. Individual values are from three independent replicates. Each replicate consisted of a pool of dissected tissues from 750-870 seedlings**.** Different letters correspond to significantly different groups (ANOVA type I, with Tukey test correction, *p-value* < 0.05).

(a)

| | Higher in RT than RR | Lower in RT than RR | Sum higher and lower expression |
|---|---|---|---|
| Only at 1µM Zn | 367 | 1129 | 1496 |
| At 1 and 200µM Zn | 959 | 3415 | 4374 |
| Only at 200µM Zn | 344 | 251 | 595 |
| **Total** | 1670 | 4795 | |

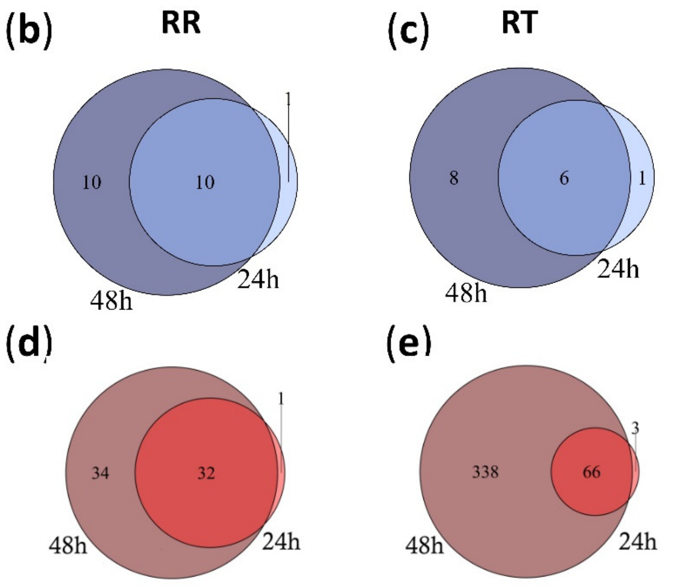


**Figure S4. Partitioning of gene expression variation between root parts or between time-points.** The experimental set-up is described in the legend of Fig. 4. **a.** Number of DEGs between RT and RR in control and/or in Zn excess conditions. **b-c.** Number of genes down-regulated in RR (**b**) and in RT (**c**) after 24h and 48h Zn excess. **d-e.** Number of genes up-regulated in RR (**d**) and in RT (**e**) after 24h and 48h Zn excess**.** Differentially expressed genes (DEG) between control and Zn excess conditions were identified separately for the 24h and 48h time points, with the following criteria: $\log_2$ (fold change) < -1 of > 1; adjusted *p-value* < 0.05.

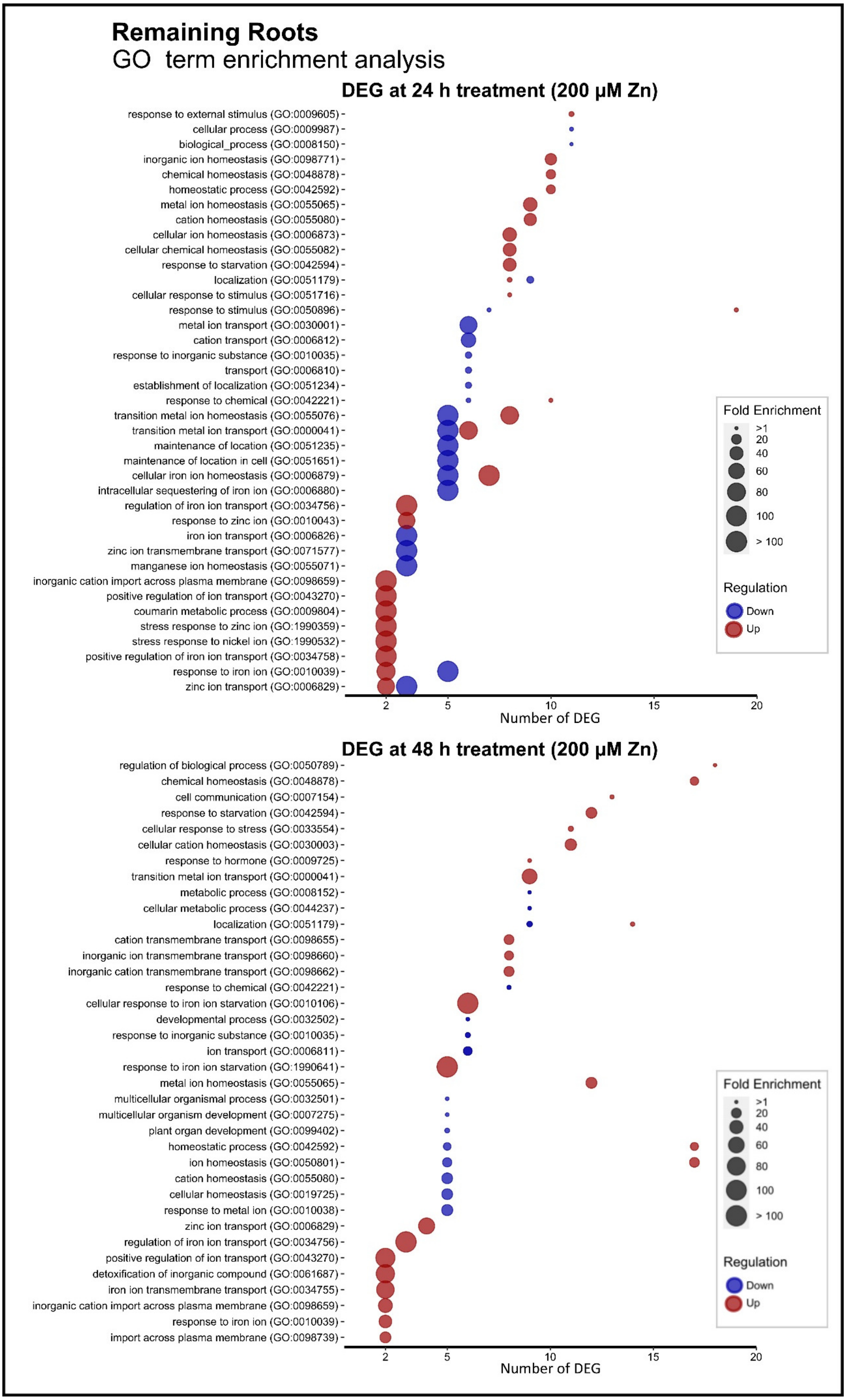
Remaining Roots
GO term enrichment analysis
DEG at 24 h treatment (200 µM Zn)
response to external stimulus (GO:0009605)
cellular process (GO:0009987)
biological_process (GO:0008150)
inorganic ion homeostasis (GO:0098771)
chemical homeostasis (GO:0048878)
homeostatic process (GO:0042592)
metal ion homeostasis (GO:0055065)
cation homeostasis (GO:0055080)
cellular ion homeostasis (GO:0006873)
cellular chemical homeostasis (GO:0055082)
response to starvation (GO:0042594)
localization (GO:0051179)
cellular response to stimulus (GO:0051716)
response to stimulus (GO:0050896)
metal ion transport (GO:0030001)
cation transport (GO:0006812)
response to inorganic substance (GO:0010035)
transport (GO:0006810)
establishment of localization (GO:0051234)
response to chemical (GO:0042221)
transition metal ion homeostasis (GO:0055076)
transition metal ion transport (GO:0000041)
maintenance of location (GO:0051235)
maintenance of location in cell (GO:0051651)
cellular iron ion homeostasis (GO:0006879)
intracellular sequestering of iron ion (GO:0006880)
regulation of iron ion transport (GO:0034756)
response to zinc ion (GO:0010043)
iron ion transport (GO:0006826)
zinc ion transmembrane transport (GO:0071577)
manganese ion homeostasis (GO:0055071)
inorganic cation import across plasma membrane (GO:0098659)
positive regulation of ion transport (GO:0043270)
coumarin metabolic process (GO:0009804)
stress response to zinc ion (GO:1990359)
stress response to nickel ion (GO:1990532)
positive regulation of iron ion transport (GO:0034758)
response to iron ion (GO:0010039)
zinc ion transport (GO:0006829)
2 5 10 15 20
Number of DEG
Fold Enrichment
>1
20
40
60
80
100
> 100
Regulation
Down
Up
DEG at 48 h treatment (200 µM Zn)
regulation of biological process (GO:0050789)
chemical homeostasis (GO:0048878)
cell communication (GO:0007154)
response to starvation (GO:0042594)
cellular response to stress (GO:0033554)
cellular cation homeostasis (GO:0030003)
response to hormone (GO:0009725)
transition metal ion transport (GO:0000041)
metabolic process (GO:0008152)
cellular metabolic process (GO:0044237)
localization (GO:0051179)
cation transmembrane transport (GO:0098655)
inorganic ion transmembrane transport (GO:0098660)
inorganic cation transmembrane transport (GO:0098662)
response to chemical (GO:0042221)
cellular response to iron ion starvation (GO:0010106)
developmental process (GO:0032502)
response to inorganic substance (GO:0010035)
ion transport (GO:0006811)
response to iron ion starvation (GO:1990641)
metal ion homeostasis (GO:0055065)
multicellular organismal process (GO:0032501)
multicellular organism development (GO:0007275)
plant organ development (GO:0099402)
homeostatic process (GO:0042592)
ion homeostasis (GO:0050801)
cation homeostasis (GO:0055080)
cellular homeostasis (GO:0019725)
response to metal ion (GO:0010038)
zinc ion transport (GO:0006829)
regulation of iron ion transport (GO:0034756)
positive regulation of ion transport (GO:0043270)
detoxification of inorganic compound (GO:0061687)
iron ion transmembrane transport (GO:0034755)
inorganic cation import across plasma membrane (GO:0098659)
response to iron ion (GO:0010039)
import across plasma membrane (GO:0098739)
2 5 10 15 20
Number of DEG
Fold Enrichment
>1
20
40
60
80
100
> 100
Regulation
Down
Up

**Figure S5. Gene Ontology (GO) enrichment analysis of genes regulated after 24h (top panel) and 48h (bottom panel) of Zn excess (200 µM Zn) in remaining roots (RR) of Arabidopsis.** The experimental set-up is described in the legend of Fig. 4. The top 15 GO for fold enrichment among genes that are up- or down-regulated by Zn excess in RR are represented in each panel. Complete data are presented in Table S4.

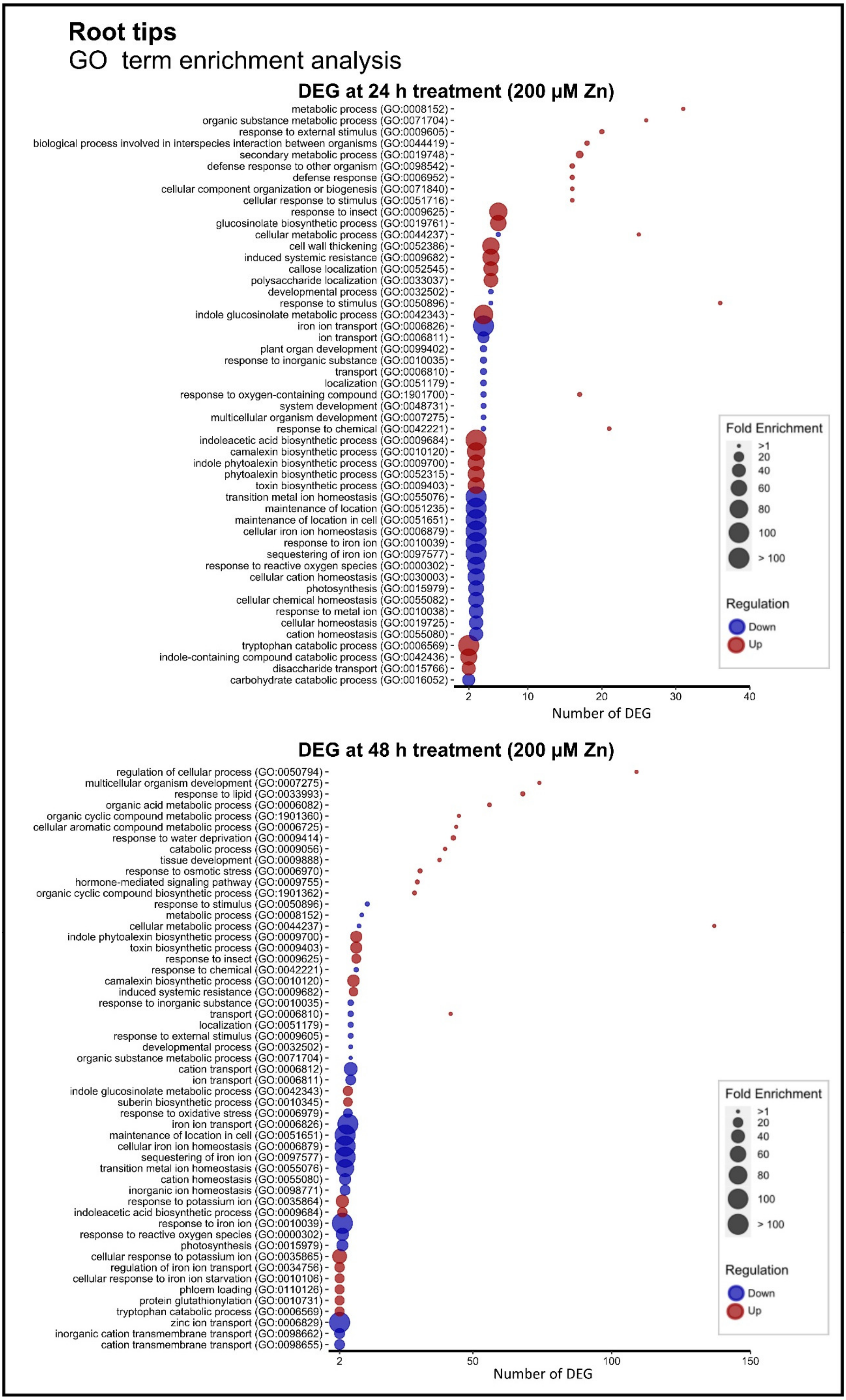
Root tips
GO term enrichment analysis
DEG at 24 h treatment (200 µM Zn)
metabolic process (GO:0008152)
organic substance metabolic process (GO:0071704)
response to external stimulus (GO:0009605)
biological process involved in interspecies interaction between organisms (GO:0044419)
secondary metabolic process (GO:0019748)
defense response to other organism (GO:0098542)
defense response (GO:0006952)
cellular component organization or biogenesis (GO:0071840)
cellular response to stimulus (GO:0051716)
response to insect (GO:0009625)
glucosinolate biosynthetic process (GO:0019761)
cellular metabolic process (GO:0044237)
cell wall thickening (GO:0052386)
induced systemic resistance (GO:0009682)
callose localization (GO:0052545)
polysaccharide localization (GO:0033037)
developmental process (GO:0032502)
response to stimulus (GO:0050896)
indole glucosinolate metabolic process (GO:0042343)
iron ion transport (GO:0006826)
ion transport (GO:0006811)
plant organ development (GO:0099402)
response to inorganic substance (GO:0010035)
transport (GO:0006810)
localization (GO:0051179)
response to oxygen-containing compound (GO:1901700)
system development (GO:0048731)
multicellular organism development (GO:0007275)
response to chemical (GO:0042221)
indoleacetic acid biosynthetic process (GO:0009684)
camalexin biosynthetic process (GO:0010120)
indole phytoalexin biosynthetic process (GO:0009700)
phytoalexin biosynthetic process (GO:0052315)
toxin biosynthetic process (GO:0009403)
transition metal ion homeostasis (GO:0055076)
maintenance of location (GO:0051235)
maintenance of location in cell (GO:0051651)
cellular iron ion homeostasis (GO:0006879)
response to iron ion (GO:0010039)
sequestering of iron ion (GO:0097577)
response to reactive oxygen species (GO:0000302)
cellular cation homeostasis (GO:0030003)
photosynthesis (GO:0015979)
cellular chemical homeostasis (GO:0055082)
response to metal ion (GO:0010038)
cellular homeostasis (GO:0019725)
cation homeostasis (GO:0055080)
tryptophan catabolic process (GO:0006569)
indole-containing compound catabolic process (GO:0042436)
disaccharide transport (GO:0015766)
carbohydrate catabolic process (GO:0016052)
2
10
20
30
40
Number of DEG
Fold Enrichment
>1
20
40
60
80
100
> 100
Regulation
Down
Up
DEG at 48 h treatment (200 µM Zn)
regulation of cellular process (GO:0050794)
multicellular organism development (GO:0007275)
response to lipid (GO:0033993)
organic acid metabolic process (GO:0006082)
organic cyclic compound metabolic process (GO:1901360)
cellular aromatic compound metabolic process (GO:0006725)
response to water deprivation (GO:0009414)
catabolic process (GO:0009056)
tissue development (GO:0009888)
response to osmotic stress (GO:0006970)
hormone-mediated signaling pathway (GO:0009755)
organic cyclic compound biosynthetic process (GO:1901362)
response to stimulus (GO:0050896)
metabolic process (GO:0008152)
cellular metabolic process (GO:0044237)
indole phytoalexin biosynthetic process (GO:0009700)
toxin biosynthetic process (GO:0009403)
response to insect (GO:0009625)
response to chemical (GO:0042221)
camalexin biosynthetic process (GO:0010120)
induced systemic resistance (GO:0009682)
response to inorganic substance (GO:0010035)
transport (GO:0006810)
localization (GO:0051179)
response to external stimulus (GO:0009605)
developmental process (GO:0032502)
organic substance metabolic process (GO:0071704)
cation transport (GO:0006812)
ion transport (GO:0006811)
indole glucosinolate metabolic process (GO:0042343)
suberin biosynthetic process (GO:0010345)
response to oxidative stress (GO:0006979)
iron ion transport (GO:0006826)
maintenance of location in cell (GO:0051651)
cellular iron ion homeostasis (GO:0006879)
sequestering of iron ion (GO:0097577)
transition metal ion homeostasis (GO:0055076)
cation homeostasis (GO:0055080)
inorganic ion homeostasis (GO:0098771)
response to potassium ion (GO:0035864)
indoleacetic acid biosynthetic process (GO:0009684)
response to iron ion (GO:0010039)
response to reactive oxygen species (GO:0000302)
photosynthesis (GO:0015979)
cellular response to potassium ion (GO:0035865)
regulation of iron ion transport (GO:0034756)
cellular response to iron ion starvation (GO:0010106)
phloem loading (GO:0110126)
protein glutathionylation (GO:0010731)
tryptophan catabolic process (GO:0006569)
zinc ion transport (GO:0006829)
inorganic cation transmembrane transport (GO:0098662)
cation transmembrane transport (GO:0098655)
2
50
100
150
Number of DEG
Fold Enrichment
>1
20
40
60
80
100
> 100
Regulation
Down
Up

**Figure S6. Gene Ontology (GO) enrichment analysis of genes regulated after 24h (top panel) and 48h (bottom panel) of Zn excess (200 µM Zn) in root tips (RT) of Arabidopsis.** The experimental set-up is described in the legend of Fig. 4. The top 15 GO for fold enrichment among genes that are up- or down-regulated by Zn excess in RT are represented in each panel. Complete data are presented in Table S4.

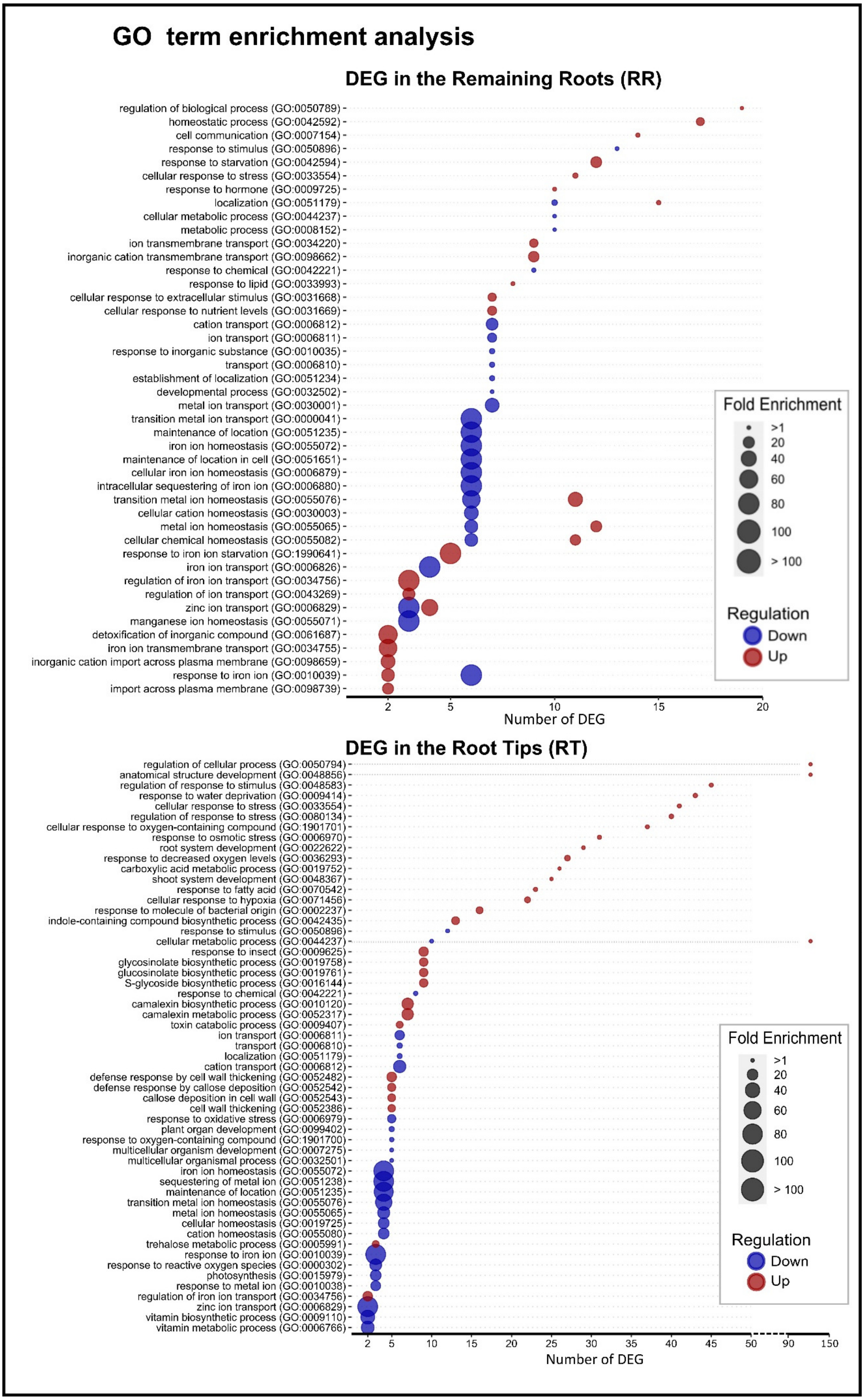

GO term enrichment analysis
DEG in the Remaining Roots (RR)
regulation of biological process (GO:0050789)
homeostatic process (GO:0042592)
cell communication (GO:0007154)
response to stimulus (GO:0050896)
response to starvation (GO:0042594)
cellular response to stress (GO:0033554)
response to hormone (GO:0009725)
localization (GO:0051179)
cellular metabolic process (GO:0044237)
metabolic process (GO:0008152)
ion transmembrane transport (GO:0034220)
inorganic cation transmembrane transport (GO:0098662)
response to chemical (GO:0042221)
response to lipid (GO:0033993)
cellular response to extracellular stimulus (GO:0031668)
cellular response to nutrient levels (GO:0031669)
cation transport (GO:0006812)
ion transport (GO:0006811)
response to inorganic substance (GO:0010035)
transport (GO:0006810)
establishment of localization (GO:0051234)
developmental process (GO:0032502)
metal ion transport (GO:0030001)
transition metal ion transport (GO:0000041)
maintenance of location (GO:0051235)
iron ion homeostasis (GO:0055072)
maintenance of location in cell (GO:0051651)
cellular iron ion homeostasis (GO:0006879)
intracellular sequestering of iron ion (GO:0006880)
transition metal ion homeostasis (GO:0055076)
cellular cation homeostasis (GO:0030003)
metal ion homeostasis (GO:0055065)
cellular chemical homeostasis (GO:0055082)
response to iron ion starvation (GO:1990641)
iron ion transport (GO:0006826)
regulation of iron ion transport (GO:0034756)
regulation of ion transport (GO:0043269)
zinc ion transport (GO:0006829)
manganese ion homeostasis (GO:0055071)
detoxification of inorganic compound (GO:0061687)
iron ion transmembrane transport (GO:0034755)
inorganic cation import across plasma membrane (GO:0098659)
response to iron ion (GO:0010039)
import across plasma membrane (GO:0098739)
2
5
10
15
20
Number of DEG
Fold Enrichment
>1
20
40
60
80
100
> 100
Regulation
Down
Up
DEG in the Root Tips (RT)
regulation of cellular process (GO:0050794)
anatomical structure development (GO:0048856)
regulation of response to stimulus (GO:0048583)
response to water deprivation (GO:0009414)
cellular response to stress (GO:0033554)
regulation of response to stress (GO:0080134)
cellular response to oxygen-containing compound (GO:1901701)
response to osmotic stress (GO:0006970)
root system development (GO:0022622)
response to decreased oxygen levels (GO:0036293)
carboxylic acid metabolic process (GO:0019752)
shoot system development (GO:0048367)
response to fatty acid (GO:0070542)
cellular response to hypoxia (GO:0071456)
response to molecule of bacterial origin (GO:0002237)
indole-containing compound biosynthetic process (GO:0042435)
response to stimulus (GO:0050896)
cellular metabolic process (GO:0044237)
response to insect (GO:0009625)
glycosinolate biosynthetic process (GO:0019758)
glucosinolate biosynthetic process (GO:0019761)
S-glycoside biosynthetic process (GO:0016144)
response to chemical (GO:0042221)
camalexin biosynthetic process (GO:0010120)
camalexin metabolic process (GO:0052317)
toxin catabolic process (GO:0009407)
ion transport (GO:0006811)
transport (GO:0006810)
localization (GO:0051179)
cation transport (GO:0006812)
defense response by cell wall thickening (GO:0052482)
defense response by callose deposition (GO:0052542)
callose deposition in cell wall (GO:0052543)
cell wall thickening (GO:0052386)
response to oxidative stress (GO:0006979)
plant organ development (GO:0099402)
response to oxygen-containing compound (GO:1901700)
multicellular organism development (GO:0007275)
multicellular organismal process (GO:0032501)
iron ion homeostasis (GO:0055072)
sequestering of metal ion (GO:0051238)
maintenance of location (GO:0051235)
transition metal ion homeostasis (GO:0055076)
metal ion homeostasis (GO:0055065)
cellular homeostasis (GO:0019725)
cation homeostasis (GO:0055080)
trehalose metabolic process (GO:0005991)
response to iron ion (GO:0010039)
response to reactive oxygen species (GO:0000302)
photosynthesis (GO:0015979)
response to metal ion (GO:0010038)
regulation of iron ion transport (GO:0034756)
zinc ion transport (GO:0006829)
vitamin biosynthetic process (GO:0009110)
vitamin metabolic process (GO:0006766)
2
5
10
15
20
25
30
35
40
45
50
90
150
Number of DEG
Fold Enrichment
>1
20
40
60
80
100
> 100
Regulation
Down
Up

**Figure S7. Gene Ontology (GO) enrichment analysis of genes regulated upon Zn excess (200 µM Zn) in remaining roots (RR, top) and root tips (RT, bottom) of Arabidopsis.** The experimental set-up is described in the legend of Fig. 4. DEGs identified after 24h and 48h of Zn excess were pooled prior analysis. The top 15 GO for fold enrichment among genes that are up- or down-regulated by Zn excess are represented in each panel. Complete data are presented in Table S4.

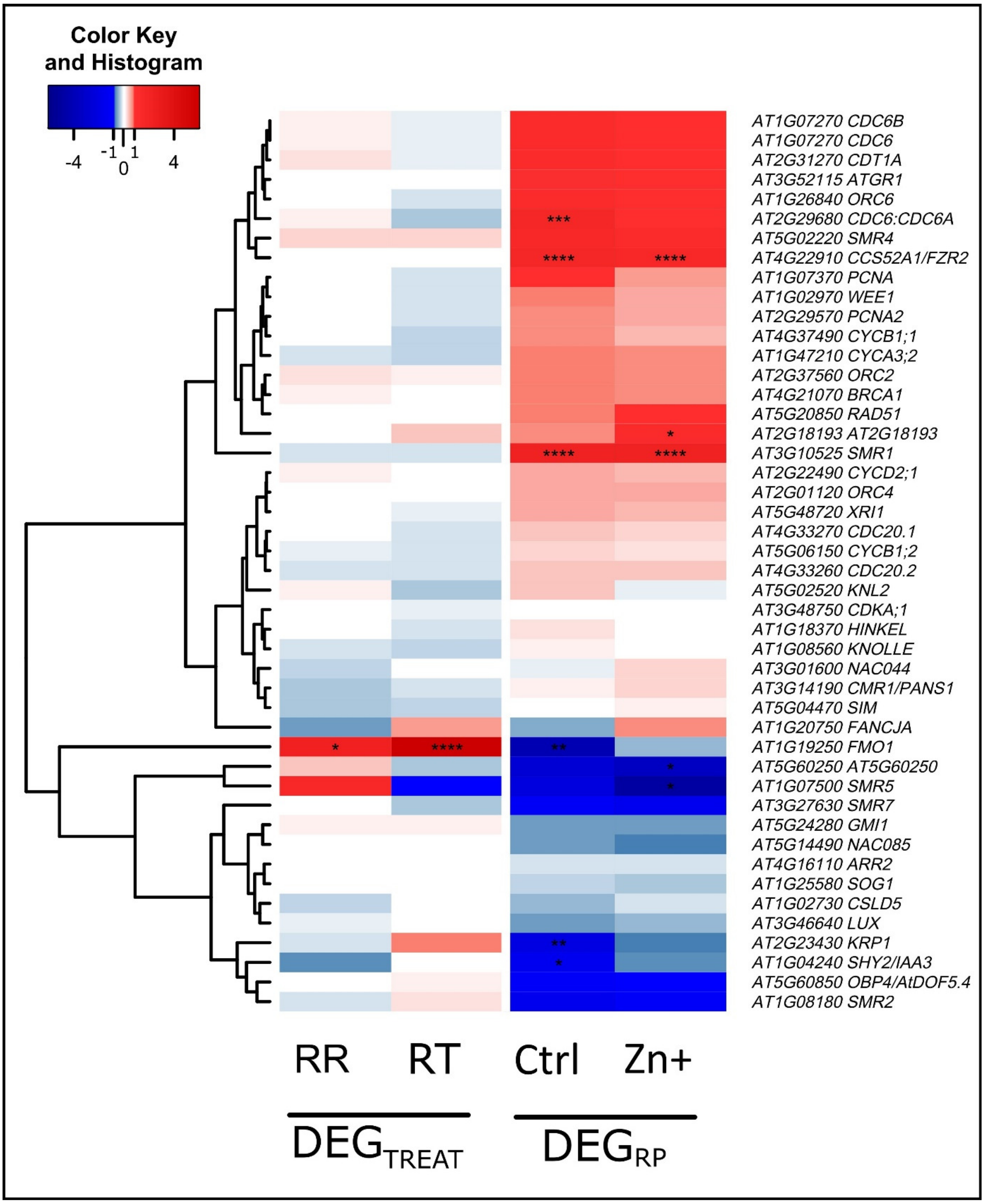


**Figure S8. Impact of Zn excess on the expression of cell cycle phase, endocycle and DNA damage marker genes in Arabidopsis roots.** Heatmap of $log_2$(fold change) of gene expression between Zn excess (Zn+) and control conditions ($DEG_{TREAT}$) or between RT and RR ($DEG_{RP}$), based on RNA-Seq data. Statistically-significant changes in gene expression (adjusted *p-value* < 0.05) are marked with stars : " * " for pval < 0.05, " ** " for pval <0.01, " *** " for pval < 0.001, " **** " for pval < 0.0001.

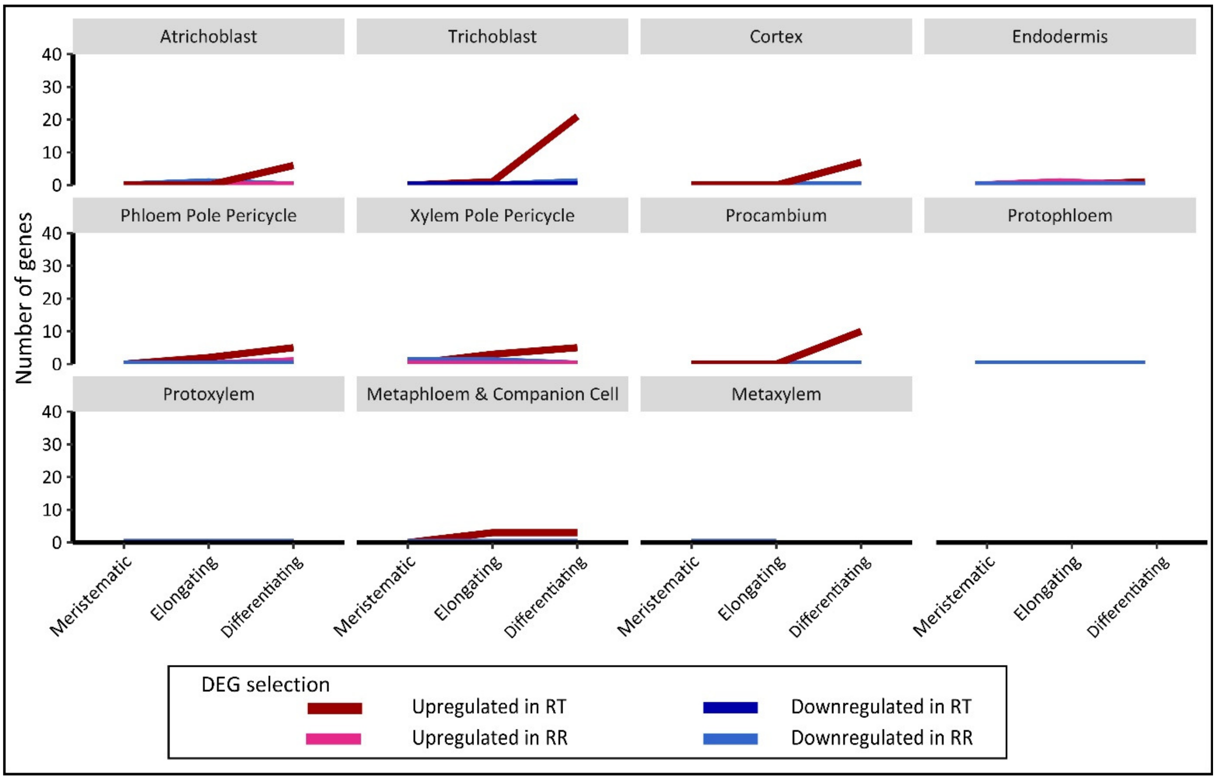


**Figure S9. Impact of Zn excess on differentiation in Arabidopsis roots.** Progression of the differentiation in the root tips upon Zn excess. Bulk RNA-Seq data from root tip (RT) and remaining root (RR) samples upon growth in control (1 µM Zn) or Zn excess (200 µM Zn, 48h) conditions were cross-referenced with the cell-type specific description of gene expression in roots provided by Shahan *et al.,* (2022). The figure shows the number of genes differentially expressed between excess and control conditions that are among the 50 genes that are specific markers for each cell line and development stage.

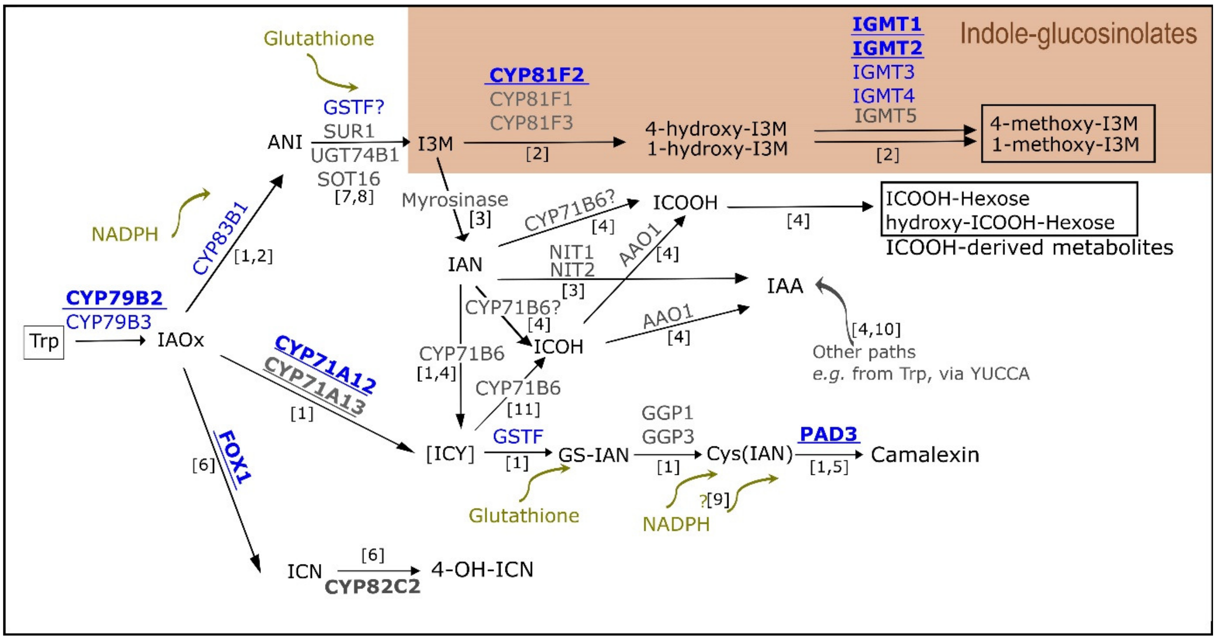


**Figure S10. The camalexin biosynthetic pathway upon Zn excess in Arabidopsis roots.** Representation of the WRKY33 regulon (camalexin, Indole glucosinolates (IG) and indole-3 carbonylnitrile (ICN) synthetic pathways from tryptophan (Trp)) and their relation to Indole-acetic acid (IAA), indole-3-carboxylicacid (ICOOH) synthesis. Genes encoding underlined bold enzymes are under WRKY33 positive regulation. Straight arrows represent one or several enzymatic reactions catalyzed by the enzymes associated to the arrow. Genes encoding blue enzymes are upregulated in root tips (RT) after 48h of Zn excess (200 µM Zn). Metabolites in black squares were identified in Figure 6. ANI : Acid Nitro indole, GS-IAN : Glutathione - indole-3-acetonitrile, IAOx : indole-3-acetaldoxime, IAN : indole-3-acetonitrile, ICN : indole-3 carbonylnitrile, ICOH : Indole-3-carbaldehyde, ICOOH : indole-3-carboxylicacid, ICY : indol-3cyanohydrin, IMG : Indole-3 methylglucosinolate, I3M : indol-3 ylmethylglucosinolate, Trp : Tryptophan.

**References:** [1] (Mucha *et al.*, 2019), [2] (Pfalz *et al.*, 2011, 2016), [3] (Hornbacher *et al.*, 2022), [4] (Böttcher *et al.*, 2014), [5] (Nafisi *et al.*, 2007),[6] (Rajniak *et al.*, 2015), [7] (Mikkelsen *et al.*, 2004),[8] (Sønderby *et al.*, 2010), [9] (Böttcher *et al.*, 2009), [10] (Kriechbaumer *et al.*, 2012), [11] (Pastorczyk *et al.*, 2020).

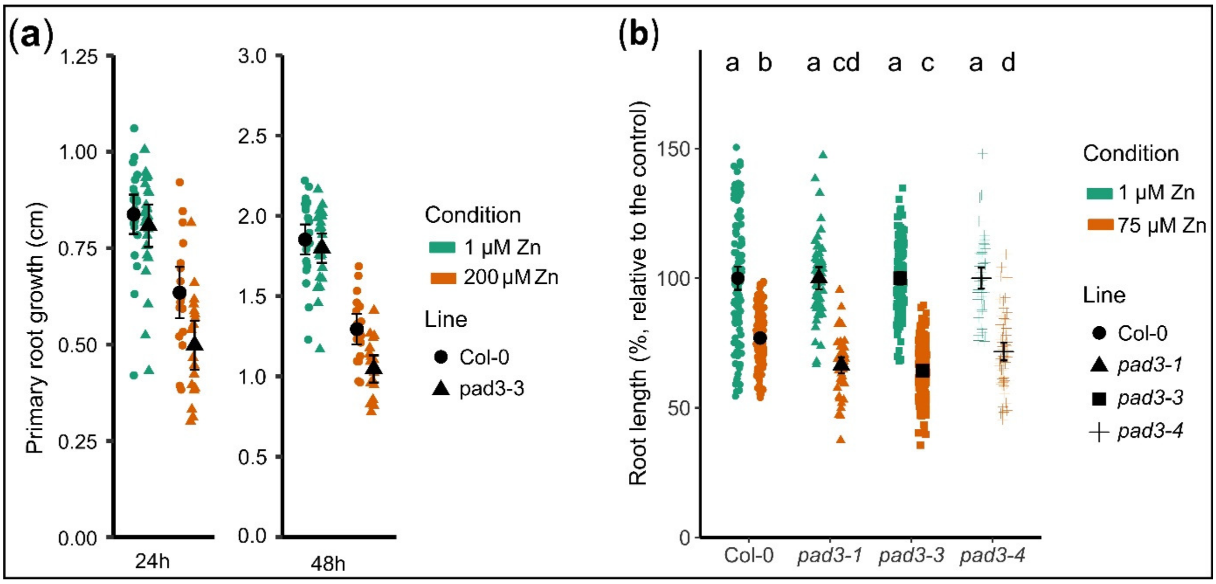


**Figure S11. Primary root growth inhibition on *pad3* knocked-out mutants. a.** Primary root growth of the *pad3-1* Arabidopsis mutant upon Zn excess. Seedlings were germinated and grown for one week in control agar plates (1 µM Zn), then transferred for another week onto new control or Zn excess (200 µM Zn) plates. Values are from 3 biological replicates, each including 9 seedlings. **b.** Col-0 and 3 different *pad3* knocked-out mutant seedlings were germinated and grown for 11 days on control (1 µM Zn) or Zn excess (75 µM Zn) agar plates. Primary root growth was measured after 11 days of growth, relative to their respective control condition, each sample included from 50 to 230 independent measurements. Different letters correspond to significantly different groups (ANOVA type I, with Tukey test correction, *p-value* < 0.05).

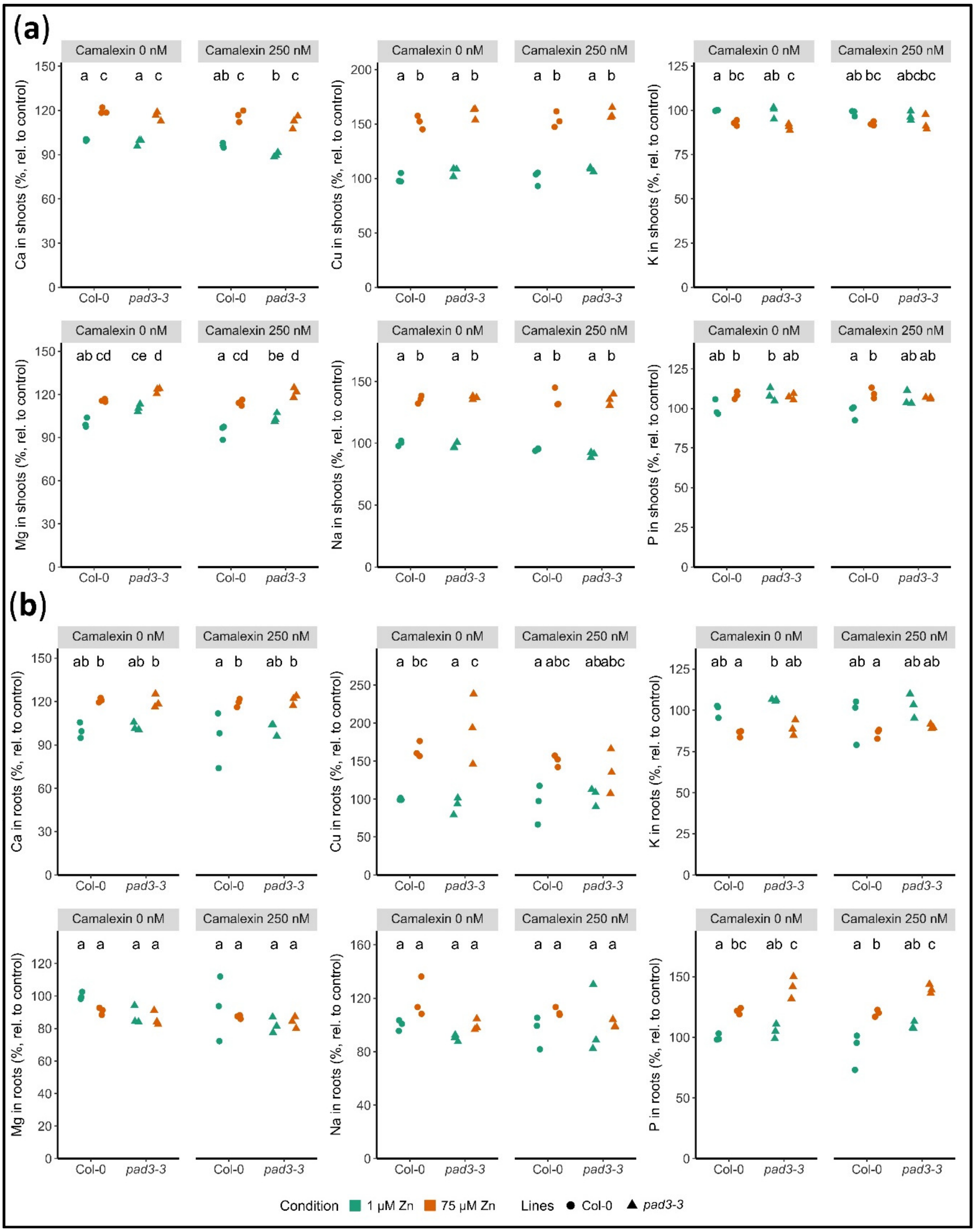


**Figure S12. Ionome profiling of the roots and shoots upon Zn excess and/or camalexin treatment in Arabidopsis.** Col-0 and *pad3-3* knocked-out mutant seedlings were germinated and grown for 11 days on control (1 µM Zn) or Zn excess (75 µM Zn) agar plates, and control (0 nM) or exogenous camalexin treatment (250 nM). Elemental concentrations are relative to their concentrations in Col-0 in control condition, from 3 biological replicates, each including 60 to 90 seedlings. Zn, Fe and Mn concentrations are presented in Fig. **7d**. Different letters correspond to significantly different groups (ANOVA type I, with Tukey test correction, *p-value* < 0.05).

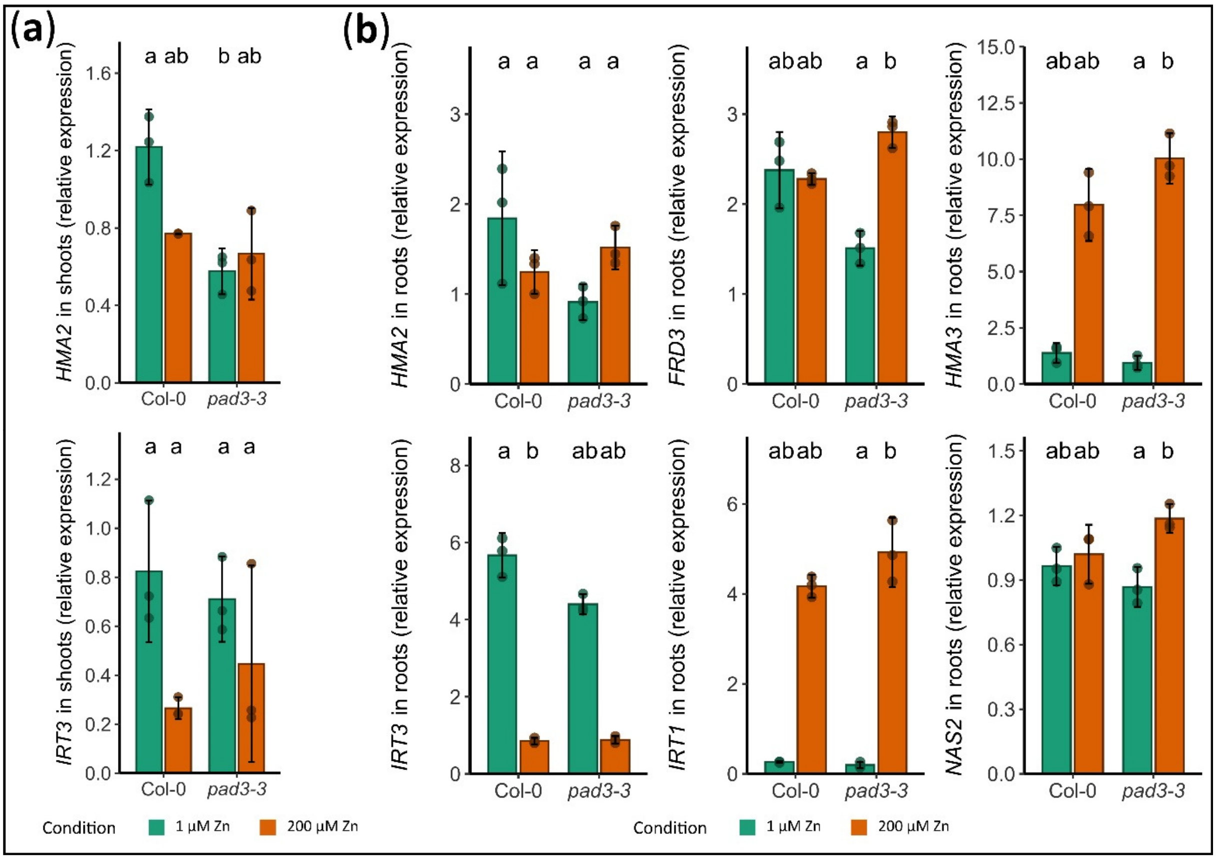


**Figure S13. Expression of zinc and iron homeostasis genes in the *pad3-3* Arabidopsis mutant.** Col-0 and *pad3-3* knocked-out mutant seedlings were germinated and grown for 11 days on control (1 µM Zn) or Zn excess (75 µM Zn) agar plates. Gene expression was assessed by RT-qPCR and normalized over 3 reference genes. Data were obtained from 3 biological replicates (40-60 seedlings/replicate), each with 3 technical replicates. Different letters correspond to significantly different groups (ANOVA type I, with Tukey test correction, *p-value* < 0.05).

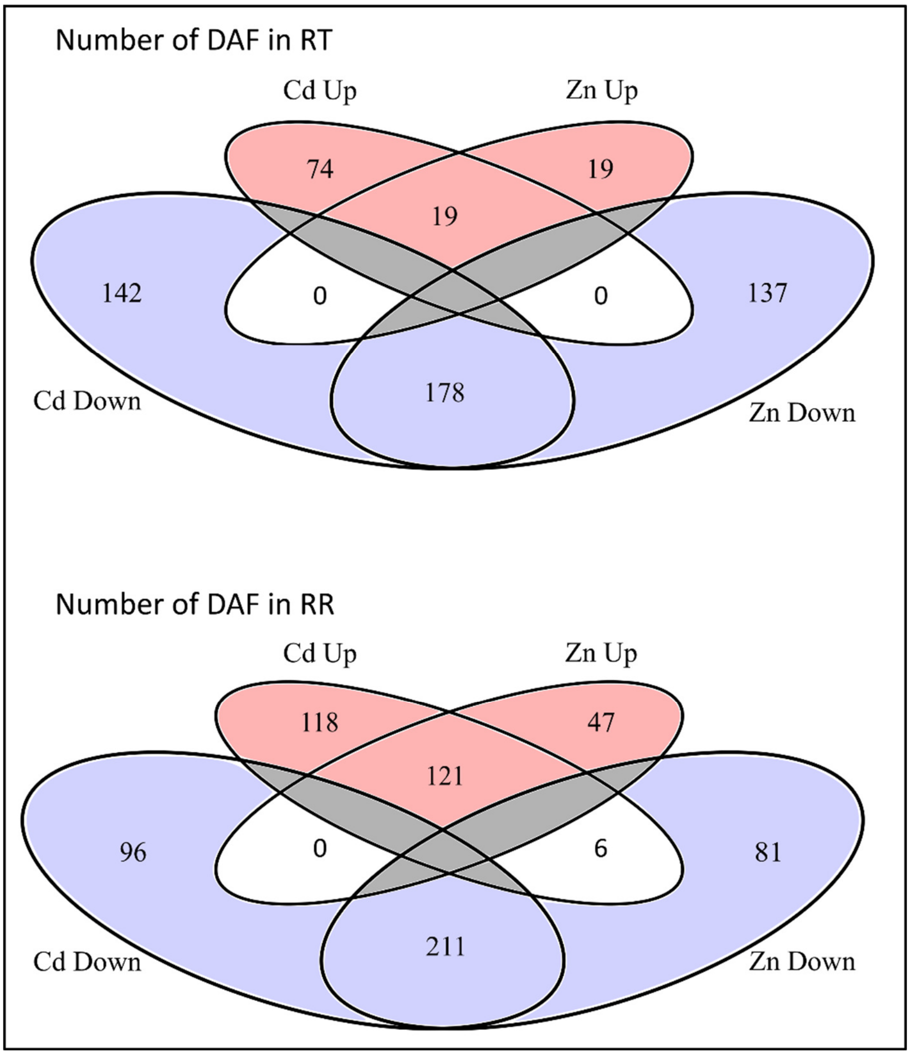


**Figure S14. Comparison of the impact of Zn excess and Cd stress on the metabolomic profile of the root tip and the remaining root system in Arabidopsis.** Seedlings were germinated and grown for one week in control agar plates (1 µM Zn), then transferred for 48h onto new control (1 µM Zn), Zn excess (200 µM Zn) or Cd (25 µM) plates. Root tip (RT) and remaining root (RR, 2-3 mm segment ~mid-length of the root) samples were then submitted to untargeted metabolomic. Data were normalized according to the total count features of the control sample, for each root part. Control, Zn excess and Cd stress experiments were simultaneously performed. The Cd stress data are described in details in (Richtmann *et al.*).

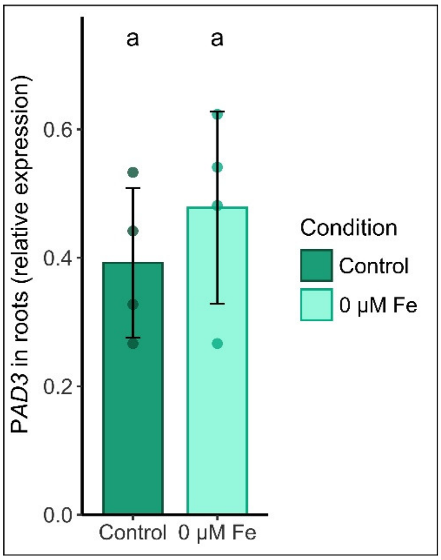


**Figure S15. *PAD3* gene expression upon Fe deficiency in Arabidopsis.** Col-0 plants were hydroponically grown for 3 weeks in control Hoagland medium, then transferred for 3 weeks in control medium (CM, 1 µM Zn, 10 µM Fe-HBED) or Fe depleted medium (1 µM Zn, 0 µM Fe). Gene expression was assessed by RT-qPCR and normalized over 2 reference genes. Data were obtained from 3 biological replicates (2-3 plants/replicate), each with 3 technical replicates. Different letters correspond to significantly different groups (ANOVA type I, with Tukey test correction, *p-value* < 0.05).

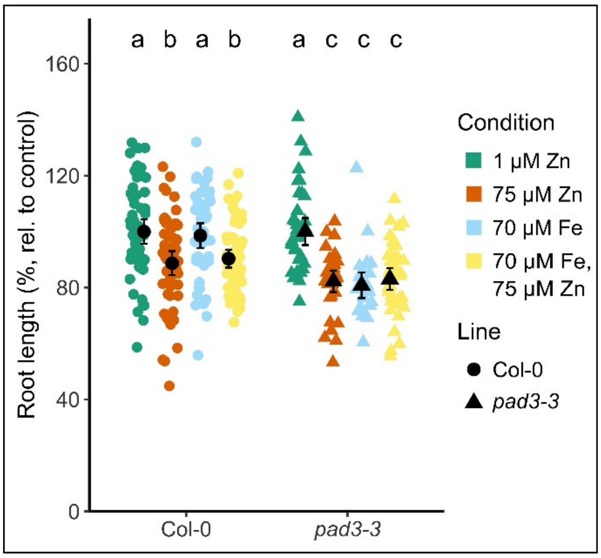


**Figure S16. Primary root growth inhibition by Zn and Fe excesses in the Arabidopsis *pad3-3* mutant.** Col-0 and *pad3-3* knocked-out mutant seedlings were germinated and grown for 11 days on control (1 µM Zn), Zn excess (75 µM Zn), Fe excess (70 µM Fe) or Fe-Zn double excess (70 µM Fe, 75 µM Zn) agar plates. Primary root growth was measured after 11 days of growth, relative to their respective control condition, each sample included 30-60 independent measurements. Different letters correspond to significantly different groups (ANOVA type I, with Tukey test correction, *p-value* < 0.05).

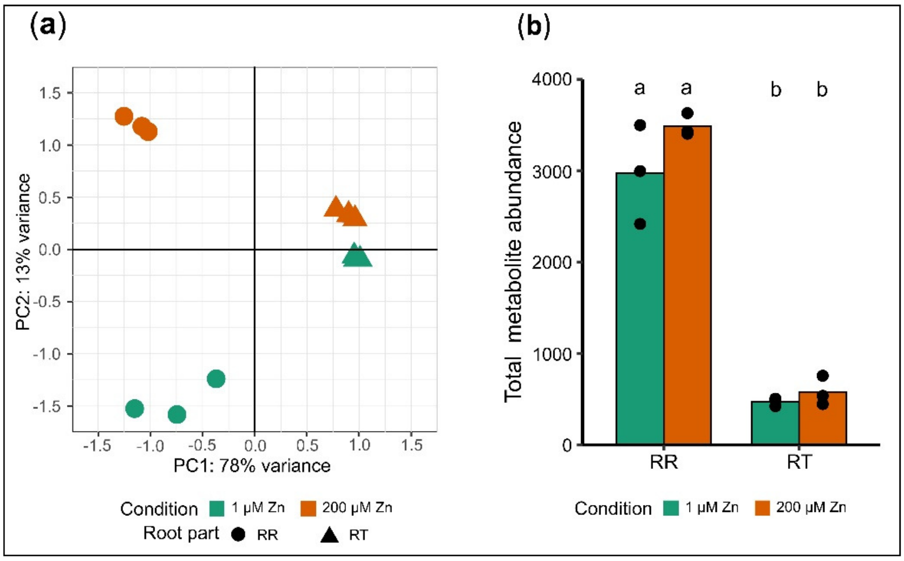


**Figure S17. Normalization of the untargeted metabolomic profiling data.** Seedlings were germinated and grown for one week in control agar plates (1 µM Zn), then transferred for 48h onto new control (1 µM Zn) or Zn excess (200 µM Zn) plates. Root tip (RT) and remaining root (RR, 2-3 mm segment ~mid-length of the root) samples were then submitted to untargeted metabolomic. **a.** Principal Component Analysis of the metabolomic data normalized according to the number of root pieces collected in each sample. **b.** Total metabolite abundance in RR and RT samples.

**Supporting Information Tables (Captions)**

The data are available as independent Exell sheets.

**Table S1.** Primers used for the RT-qPCR experiments.

**Table S2.** Genes presenting a higher or lower expression in RT than in RR, in control conditions and after 48h of Zn excess.

**Table S3.** Differentially expressed genes upon Zn excess in RT or RR, after 24h or 48h exposure to 200 µM Zn.

**Table S4.** Gene Ontology enrichment analysis amongst the differentially expressed genes upon Zn excess (24h and/or 48h) in RT and RR.

**Table S5**. Impact of Zn excess on developmental processes in the RT.

**Table S6.** Differentially expressed genes upon Zn excess annotated with nutrient stress, signaling or transport GOs.

**Table S7.** Metabolomic analysis of root samples (RR and RT) in control conditions and after 48h of Zn excess.

### References


**Anders S, Pyl PT, Huber W**. **2015**. HTSeq-A Python framework to work with high-throughput sequencing data. *Bioinformatics* **31**: 166–169.

**Bolger AM, Lohse M, Usadel B**. **2014**. Trimmomatic: A flexible trimmer for Illumina sequence data. *Bioinformatics* **30**: 2114–2120.

**Boutet S, Barreda L, Perreau F, *et al.*** 2022. Untargeted metabolomic analyses reveal the diversity and plasticity of the specialized metabolome in seeds of different *Camelina sativa* genotypes. Plant Journal **110**, 147–165.

**Böttcher C, Chapman A, Fellermeier F, Choudhary M, Scheel D, Glawischnig E**. **2014**. The biosynthetic pathway of indole-3-carbaldehyde and indole-3-carboxylic acid derivatives in arabidopsis. *Plant Physiology* **165**: 841–853.

**Böttcher C, Westphal L, Schmotz C, Prade E, Scheel D, Glawischnig E**. **2009**. The multifunctional enzyme CYP71b15 (Phytoalexin deficient3) converts cysteine-Indole-3-acetonitrile to camalexin in the Indole-3-acetonitrile metabolic network of *Arabidopsis thaliana*. *Plant Cell* **21**: 1830–1845.

**Cheng CY, Krishnakumar V, Chan AP, Thibaud-Nissen F, Schobel S, Town CD**. **2017**. Araport11: a complete reannotation of the *Arabidopsis thaliana* reference genome. *Plant Journal* **89**: 789–804.

**Dührkop K, Nothias LF, Fleischauer M, Reher R, Ludwig M, Hoffmann MA, Petras D, Gerwick WH, Rousu J, Dorrestein PC, *et al.*** **2021**. Systematic classification of unknown metabolites using high-resolution fragmentation mass spectra. *Nature Biotechnology* **39**: 462–471.

**Glazebrook J, Zook M, Mert IF, Kagan I, Rogers EE, Crute IR, Holub EB, Hammerschmidt R, Ausubelt FM**. **1997**. Phytoalexin-Deficient Mutants of Arabidopsis Reveal That <i>PAD4<\i> Encodes a Regulatory Factor and That Four <i>PAD<\i> Genes Contribute to Downy Mildew Resistance. *Genetics* **146**: 381–392.

**Di Guida R, Engel J, Allwood JW, Weber RJM, Jones MR, Sommer U, Viant MR, Dunn WB**. **2016**. Non-targeted UHPLC-MS metabolomic data processing methods: a comparative investigation of normalisation, missing value imputation, transformation and scaling. *Metabolomics : Official journal of the Metabolomic Society* **12**.

**Holmes E, Loo RL, Stamler J, Bictash M, Yap IKS, Chan Q, Ebbels T, De Iorio M, Brown IJ, Veselkov KA, *et al.*** **2008**. Human metabolic phenotype diversity and its association with diet and blood pressure. *Nature 2008 453:7193* **453**: 396–400.

**Hornbacher J, Horst-Niessen I, Herrfurth C, Feussner I, Papenbrock J**. **2022**. First experimental evidence suggests use of glucobrassicin as source of auxin in drought-stressed Arabidopsis thaliana. *Frontiers in Plant Science* **13**: 1–17.

**Kriechbaumer V, Wang P, Hawes C, Abell BM**. **2012**. Alternative splicing of the auxin biosynthesis gene YUCCA4 determines its subcellular compartmentation. *Plant Journal* **70**: 292–302.

**Love MI, Huber W, Anders S**. **2014**. Moderated estimation of fold change and dispersion for RNA-seq data with DESeq2. *Genome Biology* **15**: 1–21.

**Mikkelsen MD, Naur P, Halkier BA**. **2004**. Arabidopsis mutants in the C-S lyase of glucosinolate biosynthesis establish a critical role for indole-3-acetaldoxime in auxin homeostasis. *Plant Journal* **37**: 770–777.

**Mucha S, Heinzlmeir S, Kriechbaumer V, Strickland B, Kirchhelle C, Choudhary M, Kowalski N, Eichmann R, Hückelhoven R, Grill E, *et al.* 2019**. The Formation of a Camalexin Biosynthetic Metabolon. *The Plant cell* **31**: 2697–2710.

**Myers OD, Sumner SJ, Li S, Barnes S, Du X**. **2017**. One Step Forward for Reducing False Positive and False Negative Compound Identifications from Mass Spectrometry Metabolomics Data: New Algorithms for Constructing Extracted Ion Chromatograms and Detecting Chromatographic Peaks. *Analytical Chemistry* **89**: 8696–8703.

**Nafisi M, Goregaoker S, Botanga CJ, Glawischnig E, Olsen CE, Halkier BA, Glazebrook J**. **2007**. Arabidopsis cytochrome P450 monooxygenase 71A13 catalyzes the conversion of indole-3-acetaldoxime in camalexin synthesis. *Plant Cell* **19**: 2039–2052.

**Olivon F, Elie N, Grelier G, Roussi F, Litaudon M, Touboul D**. **2018**. MetGem Software for the Generation of Molecular Networks Based on the t-SNE Algorithm. *Analytical chemistry* **90**: 13900–13908.

**Pang Z, Zhou G, Ewald J, Chang L, Hacariz O, Basu N, Xia J**. **2022**. Using MetaboAnalyst 5.0 for LC–HRMS spectra processing, multi-omics integration and covariate adjustment of global metabolomics data. *Nature Protocols 2022 17:8* **17**: 1735–1761.

**Pastorczyk M, Kosaka A, Piślewska-Bednarek M, López G, Frerigmann H, Kułak K, Glawischnig E, Molina A, Takano Y, Bednarek P**. **2020**. The role of CYP71A12 monooxygenase in pathogen-triggered tryptophan metabolism and Arabidopsis immunity. *New Phytologist* **225**: 400–412.

**Perilli S, Sabatini S**. **2010**. Analysis of Root Meristem Size Development. *Plant Developmental Biology: Methods and Protocols* **655**: 177–187.

**Persson DP, Chen A, Aarts MGM, Salt DE, Schjoerring JK, Husted S**. **2016**. Multi-element bioimaging of *Arabidopsis thaliana* roots. *Plant Physiology* **172**: 835–847.

**Pfalz M, Mikkelsen MD, Bednarek P, Olsen CE, Halkier BA, Kroymann J**. **2011**. Metabolic engineering in *Nicotiana benthamiana* reveals key enzyme functions in Arabidopsis indole glucosinolate modification. *Plant Cell* **23**: 716–729.

**Pfalz M, Mukhaimar M, Perreau F, Kirk J, Hansen CIC, Olsen CE, Agerbirk N, Kroymann J**. **2016**. Methyl transfer in glucosinolate biosynthesis mediated by indole glucosinolate O-methyltransferase 5. *Plant Physiology* **172**: 2190–2203.

**Rajniak J, Barco B, Clay NK, Sattely ES**. **2015**. A new cyanogenic metabolite in Arabidopsis required for inducible pathogen defence. *Nature* **525**: 376–379.

**Richtmann L, Thiebaut N, Sarthou M, Ranjan A, Boutet S, Hanikenne M, Verbruggen N, Clemens S**. A multi-omics analysis of Arabidopsis thaliana root tips under Cd exposure: Insights into HY5's role in limiting accumulation. *New Phytologist*.

**Shahan R, Hsu CW, Nolan TM, *et al.*** 2022. A single-cell Arabidopsis root atlas reveals developmental trajectories in wild-type and cell identity mutants. Developmental Cell **57**, 543-560.e9

**Sønderby IE, Geu-Flores F, Halkier BA**. **2010**. Biosynthesis of glucosinolates - gene discovery and beyond. *Trends in Plant Science* **15**: 283–290.

**Tewes LJ, Stolpe C, Kerim A, Krämer U, Müller C**. **2018**. Metal hyperaccumulation in the Brassicaceae species *Arabidopsis halleri* reduces camalexin induction after fungal pathogen attack. *Environmental and Experimental Botany* **153**: 120–126.
